# Kilobase-scale compartments enabled by CRUSH reveal regulatory programs across cell types, single-cells, and ancient mammoths

**DOI:** 10.64898/2026.04.25.719469

**Authors:** Achyuth Kalluchi, Ben Nolan, Timothy E. Reznicek, Hannah L. Harris, Devika Udupa, Christopher T. Cummings, M. Jordan Rowley

## Abstract

How chromatin is spatially organized in the nucleus has long been studied through the lens of large-scale A/B compartments, but whether these sizes reflect true biological units or analytical artifacts has remained unclear. We find that limits imposed by the conventional eigenvector-based compartment calling have required extreme sequencing depth and coarse resolution, obscuring regulatory-scale organization. We developed CRUSH to iteratively refine compartments to 1 kb resolution without the need for extreme sequencing depth, we show that kilobase-scale A/B segregation (micro-compartments) is evident across cell types. Across these maps, we demonstrate that RNA polymerase II pausing contributes to a sub-genic compartment signature at the transcription start site and that active enhancers almost universally occupy the A compartment. We then show that fine-scale compartment maps can resolve cancer subtype-specific regulatory programs, single-cell tissue identity, and cold-adaptation regulomes in a 52,000-year-old woolly mammoth. These findings establish chromatin compartmentalization as a gene-scale regulatory feature with broad implications for development, disease, and genome evolution.

**Graphical Abstract:** 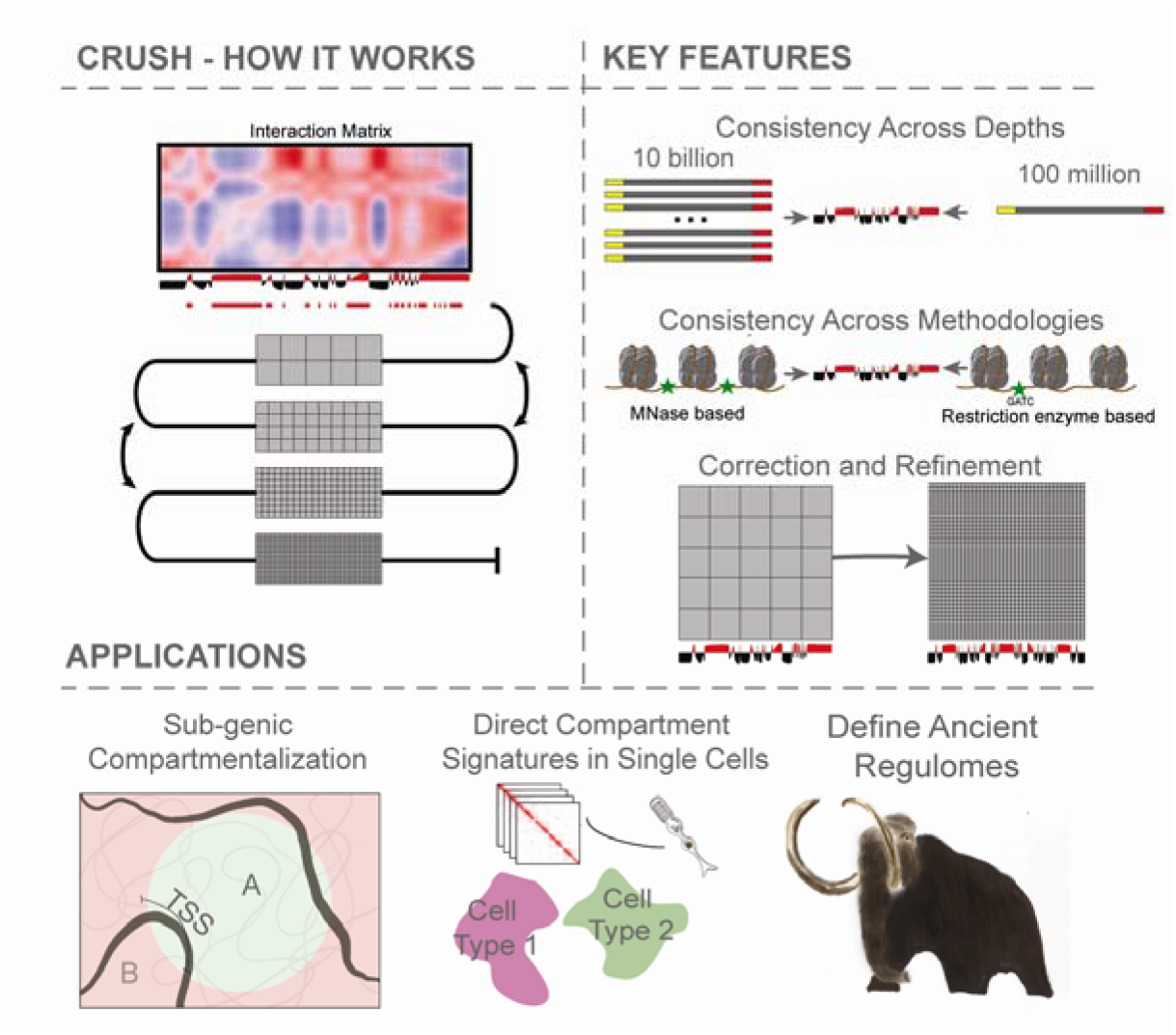

CRUSH uses iterative resolution walking to identify and correct A/B compartment measurements. This allows identification of A/B compartments at kilobase resolution with low sequencing depth and consistency across methodologies. Kilobase-scale compartments reveal sub-genic compartment organization, compartment identities in single-cells, and a distinct cold-adapted regulome in mammoths.

## Main Text

The three-dimensional organization of the genome plays a fundamental role in gene regulation, with chromatin partitioned into active (A) and inactive (B) compartments that reflect distinct nuclear environments (*1*). Since their discovery, compartments have been modeled as large, megabase-scale structures, an assumption that traces directly to the megabase resolution of early Hi-C maps rather than to any intrinsic biological property (*1, 2*). Recent work has challenged this view, demonstrating that compartments can be resolved to the kilobase scale, where they subdivide individual gene bodies and demarcate single regulatory elements (*3, 4*). At this resolution, compartmentalization emerges as a bottom-up property of local chromatin state rather than a top-down architectural constraint, with large compartments representing the cumulative alignment of many small, locally defined units rather than primary organizational features (*2, 3, 5*).

Resolving compartments at this scale has remained technically prohibitive (*3, 6*). Eigenvector-based methods, the field standard, prioritize global chromosomal variance, suppressing local heterogeneity and requiring extreme sequencing depth to approach kilobase resolution (*6*). These barriers have confined fine-scale compartment analysis to a handful of ultra-deeply sequenced datasets (*3, 4*), preventing systematic investigation across diverse cell types, disease states, or organisms.

Here, we introduce CRUSH (Compartment Refinement for Ultraprecise Stratification of Hi-C), which achieves kilobase-scale compartment analysis at low sequencing depths by directly measuring interaction preferences and iteratively refining calls across resolutions (see Graphical Abstract). We apply CRUSH to demonstrate that RNA polymerase II pausing contributes to a sub-genic compartment signature at transcription start sites and that A compartment localization is a near universal feature of active enhancers. We also demonstrate that cell types carry unique compartment signatures detectable in individual cells without aggregation, and that the regulatory landscape of a 52,000-year-old woolly mammoth encodes the transcription factor programs underlying cold adaptation.

### Kilobase-scale micro-compartments are detectable across cell types, methodologies, and sequencing depths

Kilobase-scale A/B compartment segregation has been reported in ultra-deeply sequenced Hi-C maps and region-capture Micro-C datasets (*3, 4*), but whether this fine-scale organization is a general feature of the genome, or is accessible in standard datasets across diverse methodologies, has not been established. To address this, we developed CRUSH, which directly measures each locus’s interaction preferences with A and B type nuclear environments across resolutions, iteratively refining compartment calls (see Supplemental Materials and Methods).

We first benchmarked CRUSH against the eigenvector in an ultra-depth Hi-C map of lymphoblastoid cells (LCLs) with 20.3 billion intrachromosomal contacts (*3*), finding high concordance in compartment calls at matched resolutions (Fig. 1A, S1A). Critically, CRUSH reproduced these compartments with only 100 million contacts, 50-fold fewer, while the eigenvector required at least 5 billion to achieve comparable assignments (Fig. 1A, S1B-D). To test methodological impacts, we next compared Hi-C and Micro-C maps from H1 cells (*7*). The eigenvector failed to produce coherent A/B assignments on some chromosomes at 1 kb and visibly noisier assignments in Micro-C at 25 kb (Fig. 1B), resulting in larger differences between methodologies than between biologically distinct cell types (Fig. 1C, S1E). CRUSH virtually eliminated this methodological bias, recovering consistent compartment calls across library types even at 1 kb, with cell-type differences exceeding methodological differences (Fig. 1C, S1E).

**Fig. 1.**
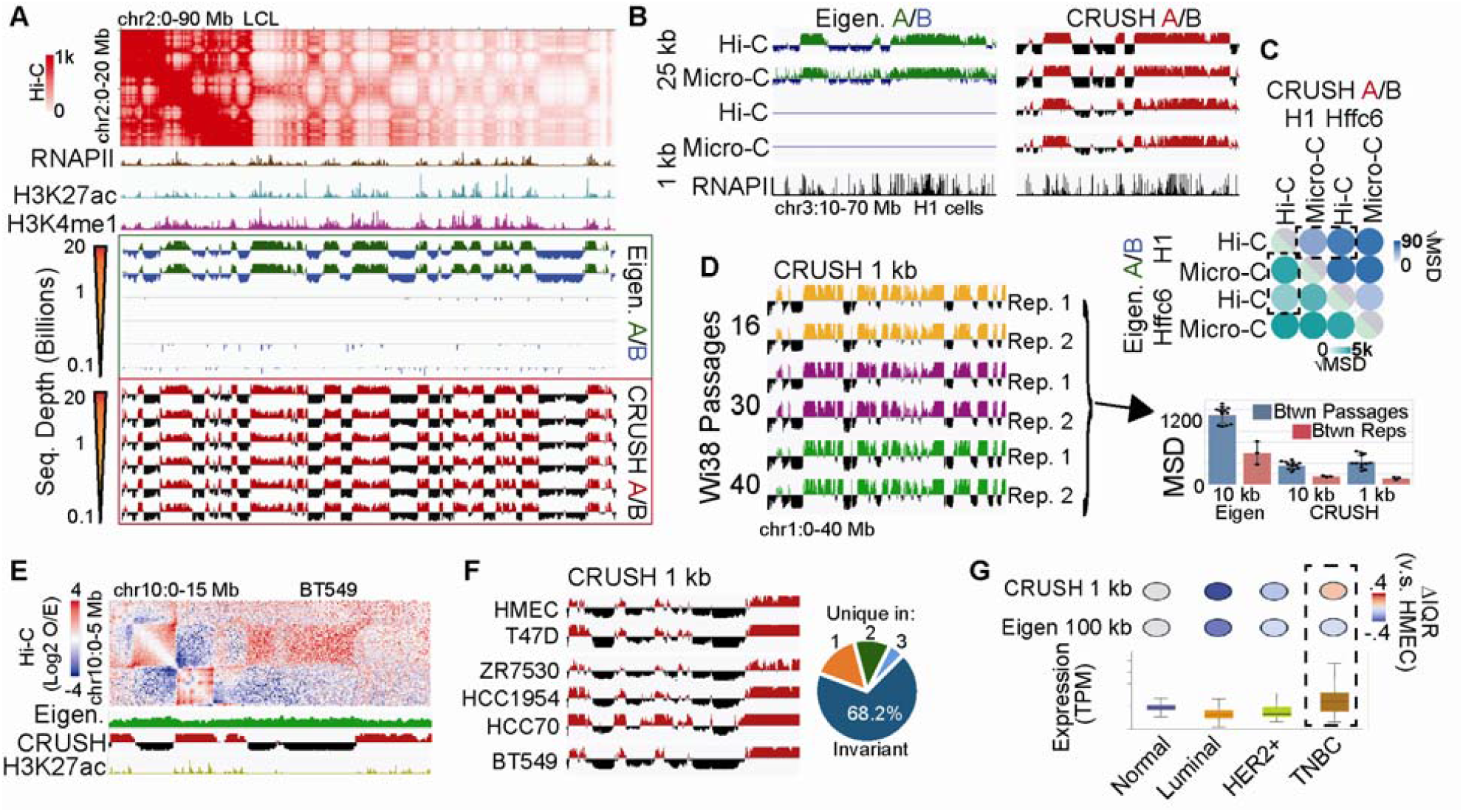
Kilobase-scale chromatin compartments are reproducible across methodologies and recoverable at low sequencing depth. **(A)** Comparison of CRUSH (red/black) and eigenvector (green/blue) defined chromatin compartments at high and low Hi-C contact depths in lymphoblastoid cell lines (LCLs). RNA Polymerase II (RNAPII), H3K27ac, and H3K4me1 are shown as representative active marks. **(B)** Compartments defined by the eigenvector (left) v.s. CRUSH (right) in H1 cells. 1 kb eigenvector tracks are empty due to a complete failure to denote coherent compartments on this chromosome. RNAPII ChIP-seq signal is shown to denote active chromatin. **(C)** Mean square difference (MSD) of CRUSH (left) and eigenvector (right) compartments between Hi-C vs. Micro-C and between H1 vs. HFFc6. Dashed boxes represent comparison between methodologies and between cell lines. Note the overall different scales. **(D)** Left: CRUSH signal at 1 kb between replicates and passages in Wi38 cells. Right: MSD comparison of CRUSH and eigenvector-derived compartment signal between passages (blue) and between replicates (red). 10 kb is shown for both eigenvector and CRUSH, whereas 1 kb is only shown for CRUSH as the eigenvector completely failed to provide coherent A/B calls for this resolution. Error bars indicate standard deviation, with individual data points shown. **(E)** Compartment intervals in BT549 cells that were missed by the eigenvector but detected by CRUSH. **(F)** Comparison of compartments at 1 kb resolution between HMEC (normal breast epithelia), T47D (luminal A), ZR7530 (luminal B), HCC1954 (HER2+), HCC70 (TNBC A), and BT549 (TNBC B) cells. Pie represents the number of compartment bins called uniformly across cell lines (invariant), or were different in one, two, three cell lines. **(G)** Expression of R5U1 in TCGA data (boxplots), and the activity status in corresponding cell lines according to either CRUSH at 1 kb, or the eigenvector (100 kb as was used previously). Box highlights an agreement identified by CRUSH that was missed by the eigenvector.

To assess sensitivity to subtler technical variation, we applied CRUSH and the eigenvector to Hi-C maps from Wi38 cells across biological replicates and cell culture passages (*8*). Passage-to-passage differences reflect a combination of actual biological change as well as technical noise, so a more precise method should reduce replicate variance while preserving biologically meaningful passage differences. CRUSH minimized inter-replicate variation while maintaining a high passage-to-replicate difference ratio (Fig. 1D). The eigenvector failed to produce coherent compartment calls at 1 kb entirely and exhibited higher inter-replicate noise at 10 kb (Fig 1D), noise that obscured biologically relevant GO terms associated with passage-dependent compartment changes, which CRUSH recovered (Fig. S1F-G).

### Correcting compartment misassignments reveals disease- and lineage-specific chromatin programs

A known limitation of eigenvector-based compartment calling is that the first principal component does not always capture the A/B compartment axis when other sources of genomic variance, such as chromosome-arm-level organization, dominate the signal and mask the underlying compartment pattern (*6*) (see Supplemental Text). For example, in Hi-C maps of *G. gallus* erythroblasts (*9*), the eigenvector reports chromosome arm separation rather than the A/B checkerboard pattern that is both visually evident in the contact map and correctly identified by CRUSH (Fig. S2A). Resolving this misassignment improved concordance between compartment calls and gene expression and revealed stage-specific compartment dynamics with functional relevance to erythroid maturation (Fig. S2B-H).

In a Hi-C map of BT549 triple-negative breast cancer (TNBC) cells (*10*), the eigenvector missed compartment intervals that are visually evident in the contact map. CRUSH identified these correctly and enabled 1 kb compartment analysis across six breast cancer cell lines (*10*) (Fig. 1E-F, S3A-B). Corrected compartment calls in T47D and BT549 showed improved concordance with H3K27ac occupancy even at 100 kb (Fig. S3C). Furthermore, compartment differences between normal breast epithelial cells (HMEC) and BT549 corresponded to H3K27ac differences at the same loci (Fig. S3D), were enriched for cancer-relevant GO terms (Fig. S3E), and matched gene expression differences observed in TCGA breast cancer patient data (*11, 12*) (Fig. 1G, S3F).

Together, these results show that eigenvector misassignments produce systematic errors that obscure real biology, erase lineage-specific compartment transitions in erythroid differentiation, and mask subtype-specific regulatory programs in breast cancer. Correcting these assignments reveals that kilobase-scale compartment organization is a faithful readout of the underlying gene regulatory state.

### RNAPII pausing creates a sub-genic compartment signature at the transcription start site

Even in the most deeply sequenced Hi-C maps, eigenvector-based compartment calls can misassign individual loci where local interaction patterns diverge from the chromosome-wide variance structure (see Supplemental Text). We asked if CRUSH, which measures each locus’ direct interaction preferences with A- and B-type environments rather than its correlation with neighbors, can correct these locus-specific errors. In the 20.3 billion contact LCL map (*3*), the majority of loci agree between methods, but at sites of disagreement CRUSH assignments better reflect both the contact map geometry and the underlying chromatin activity state (Fig. 2A-B, S4A-B).

**Fig. 2.**
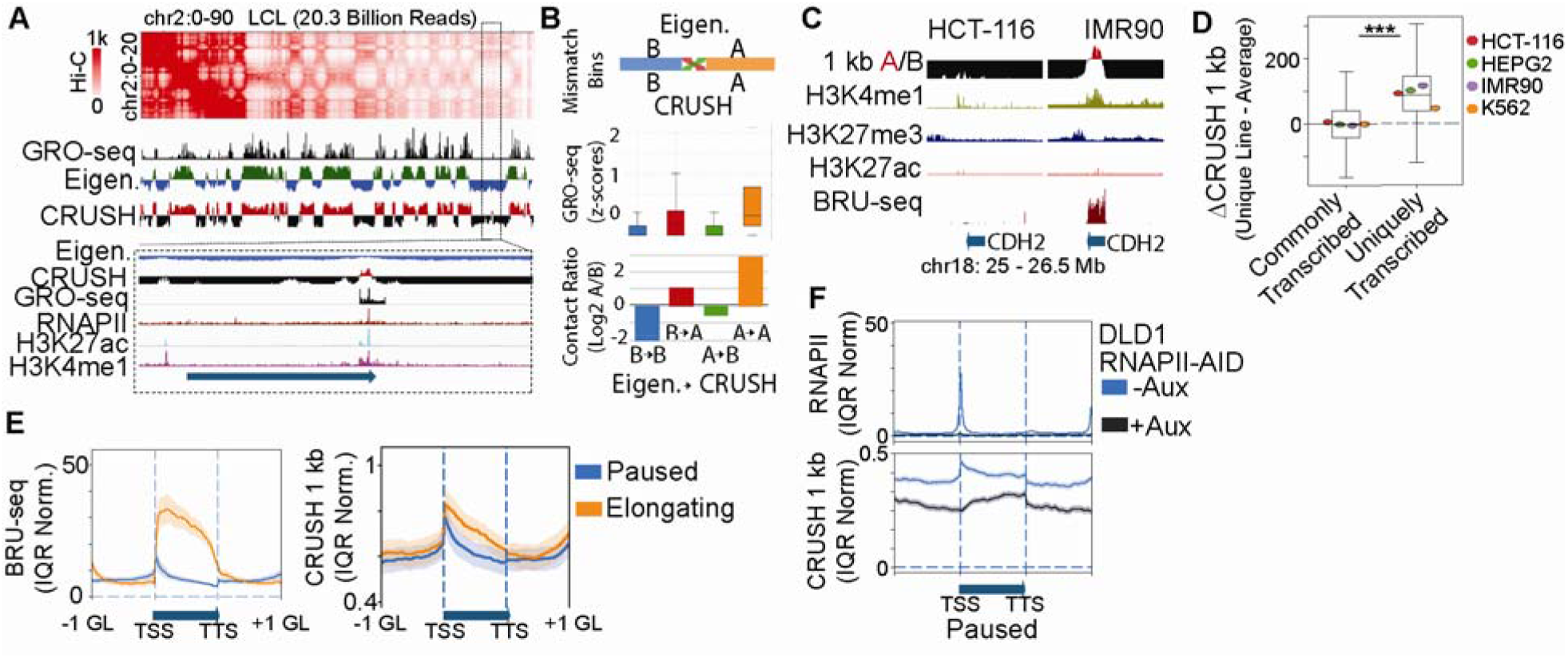
RNAPII pausing defines a sub-genic compartment signature at the TSS across cell types. **(A)** Example of a compartment call mismatch between the eigenvector and CRUSH in the deeply sequencing LCL Hi-C map. GRO-seq, RNAPII, H3K27ac, and H3K4me1 as shown to denote 1D chromatin activity state. Box denotes the zoomed-in area. **(B)** Top: Diagram proportional to the number of matching v.s. mismatched compartment calls between the eigenvector and CRUSH. Middle: GRO-seq signal and Bottom: the relative contact ratio in bins called either B (blue) or A (orange) by both algorithms, compared to those flipped B to A (red) or A to B (green) between eigenvector and CRUSH. **(C)** Example of a gene with opposite compartment association between HCT-116 and IMR90. H3k4me1, H3K27me3, H3K27ac, and BRU-seq are shown for reference. **(D)** Boxplots show differences in CRUSH scores across cell lines (ΔCRUSH) for bins that are transcribed in all four cell lines and those uniquely transcribed in one line. Wilcoxon rank-sum test, *** p < 0.001. **(E)** Average BRU-seq (left) and CRUSH score (right) profiles across gene bodies for paused (blue) and elongating (orange) genes, going −1 gene length (GL) before the TSS to +1 GL after the TTS. Profiles are shown as IQR-normalized means across all gene status compartment scores respectively across cells lines with BRU-seq data (HCT-116, HepG2, IMR90, and K562). Shaded bands represent standard deviation. **(F)** RNAPII occupancy (top) and CRUSH scores (bottom) in DLD1 RNAPII-AID cells in -auxin (blue) and +auxin (grey) conditions. Shaded bands represent standard deviation.

To characterize chromatin features that putatively drive fine-scale compartment identity genome-wide, we applied CRUSH at 1 kb across seven ENCODE cell-line Hi-C maps (A549, HCT116, HeLa, HepG2, HUVEC, IMR90, and K562) (Table S1) (*13*–*17*). A compartment bins were strongly enriched for active chromHMM states (*18*), and A compartment scores increased with the number of co-occurring active marks (Fig. S4C-D) (Table S2). H3K27me3-marked bins showed weaker compartment scores overall, but in K562 cells, where we see H3K27me3 extensively overlaps active marks, these bins were frequently assigned to the A compartment (Fig. S4E-F). This suggests that compartment status may be superseded by active marks. Indeed, examining H3K27me3 with and without active marks across the seven cell lines, we see that A compartment affinity scales with the number of co-occurring active modifications (Fig. S4G).

We then found that nascent transcription measured by BRU-seq’s ability to predict the A compartment improved at 1 kb resolution, and that transcriptional differences between cell lines corresponded directly to compartment differences (Fig. 2C-D, S5A-F). Prior work in ultra-deeply sequenced LCLs reported that paused genes, those with RNAPII stalled near the promoter, showed a spike of A compartment signal at the transcription start site (TSS) that drops sharply into the gene body (*3*). However, whether this sub-genic compartment pattern is a general feature conserved across cell types, or an idiosyncrasy of that single dataset, was unknown. Applying CRUSH across cell lines that have available BRU-seq data (HCT116, HepG2, IMR90, and K562 cells), we find that this TSS-specific A compartment signature is a consistent feature of paused genes in every cell line examined, while elongating genes maintain higher A compartment association across the full gene body (Fig. 2E, S6A).

To test whether RNAPII occupancy is causally required for this sub-genic compartment structure, we examined DLD1 cells carrying an auxin-inducible degron tag on endogenous RNAPII, a system in which RNAPII can be acutely depleted (*19*). Under basal conditions, the TSS-specific A compartment spike co-localizes with RNAPII occupancy (Fig. 2F, blue). Upon auxin-induced RNAPII degradation, this signal is specifically lost at TSSs, with the site of most loss corresponding to the highest prior RNAPII occupancy (Fig. 2F, grey; S6B). Importantly, genes do not fully convert to the B compartment upon RNAPII loss, consistent with prior observations at lower resolution (*19, 20*) and indicating that polymerase occupancy is likely only one of multiple determinants of compartment identity (see Fig. S4D). This is consistent with liquid chromatin Hi-C experiments showing that some amount of compartment identity has intrinsic physical stability (*21*). Together, these results demonstrate that RNAPII pausing at the TSS is both a marker and a mechanistic contributor to a sub-genic A compartment signature, a fine-scale organizational feature that is conserved across cell types directly coupled to the transcriptional machinery.

### Cell-type identity is encoded in single-cell compartment signatures

Single-cell Hi-C methods capture chromatin contacts from individual cells (*22*–*24*), but the sparse coverage of each map, typically orders of magnitude fewer contacts than bulk experiments, has made direct compartment calling intractable. Existing approaches instead have had to create clever work-arounds by relying on pseudo-bulk aggregation or indirect inference methods that sacrifice single-cell resolution to achieve reliable compartment assignments (*23, 25, 26*). Because CRUSH operates effectively at low contact depth, we asked whether compartments could be called directly from individual single-cell maps and whether the resulting per-cell compartment profiles carry sufficient information to identify cell type without aggregation (Fig. 3A).

**Fig. 3.**
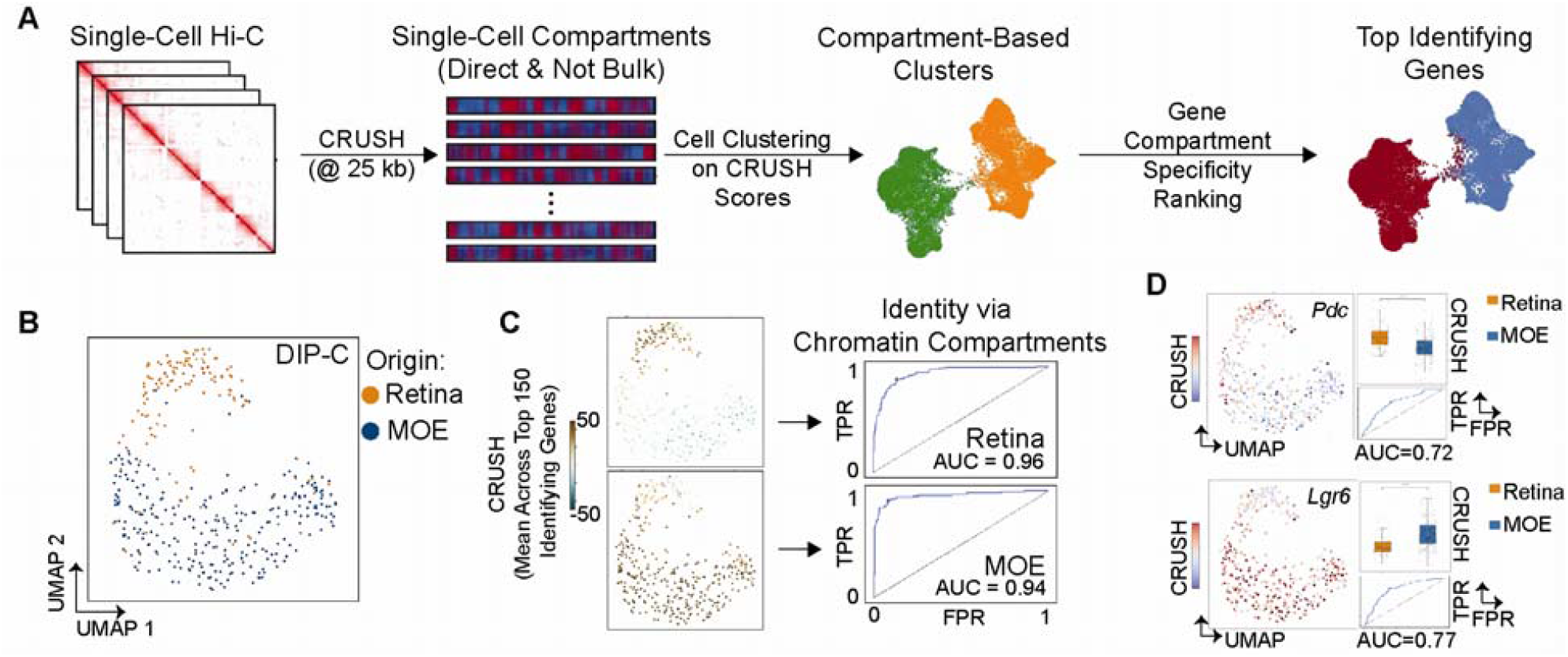
Individual cells carry stable, cell-type-specific compartment signatures identifiable in single-cell Hi-C maps. **(A)** Overview of using CRUSH for single-cell compartment analysis at 25 kb resolution to perform compartment-based clustering and the identification of gene-compartment signatures unique to each cell population. **(B)** DIP-C data comprised of retina (orange) and MOE (blue) with the UMAP based on 25 kb CRUSH called directly on each individual cell. **(C)** Left: UMAP colored by top 150 genes with unique A compartment signatures in the retina (top) and MOE (bottom). Right: Receiver Operating Characteristic (ROC) based on the True Positive Rate (TPR) and False Positive Rate (FPR) for the top 150 compartment signatures. AUC = Area Under Curve. **(D)** Left: UMAP colored by the compartment signature of *Pdc* (top) and *Lgr6* (bottom). Right: The difference in CRUSH scores between retina and MOE (boxplot) and ROC for each gene-compartment signature. *** p<.001, Wilcoxon Rank Sum test.

We first applied this approach to DIP-C data from mouse retina and main olfactory epithelium (MOE) (*24*) (Table S3). Per-cell compartment profiles computed by CRUSH robustly separated the two populations across resolutions, with optimal segregation at 25 kb (Fig. 3B, S7A-B). The top 150 differentially compartmentalized genes defined a gene-compartment signature that distinguished retina from MOE with high accuracy (Fig. 3C, Table S4-S5), outperforming eigenvector-based profiles at matched resolution and improving further at 25 kb relative to 500 kb (Fig. S7C-D). Even individual gene-compartment signatures, including *Pdc*, which is selectively active in retina, and *Lgr6*, which is selectively active in MOE, were resolved with sufficient precision to serve as single-gene classifiers (Fig. 3D, S7E-F).

To test this approach in a more challenging context, we reanalyzed an early DIP-C dataset of 33 mixed peripheral blood mononuclear cells (PBMCs) and lymphoblastoid cells (GM12878) (*23*). Here, 25 kb resolution was essential, as coarser compartment profiles completely failed to separate the two populations (Fig. S8, Table S6-S7). Together, these results establish that individual cells carry a stable, cell-type-specific compartment identity that persists against a background of single-cell heterogeneity, and that this identity is recoverable directly from sparse single-cell contact maps at kilobase-scale resolution.

### Cold-adaptation regulomes are preserved in the 52,000-year-old woolly mammoth genome

Prior work in ultra-deeply sequenced LCLs found that nearly all active enhancers localize to the A compartment at kilobase resolution (*3*), but it was unclear whether this reflected a universal organizational principle or a cell-type-specific observation. Applying CRUSH across seven ENCODE cell lines, we find that A compartment localization is a near-universal property of enhancerDB (*27*) annotated active enhancers in every cell type examined, and that this association strengthens further when enhancers are more stringently filtered for cell-type-specific H3K4me1 and H3K27ac occupancy (Fig. 4A). Enhancers unique to each cell line carry correspondingly unique A compartment signatures (Fig. 4B, S9A), and the transcription factor motifs enriched at these cell-line-specific compartment-enhancer signatures accurately reflect known cell-type biology (Fig. S9B, Table S8). KLF1 motif, for instance, dominates the K562-specific signature, consistent with their established role in erythroid specification (*28*) (Fig. 4C).

**Fig. 4.**
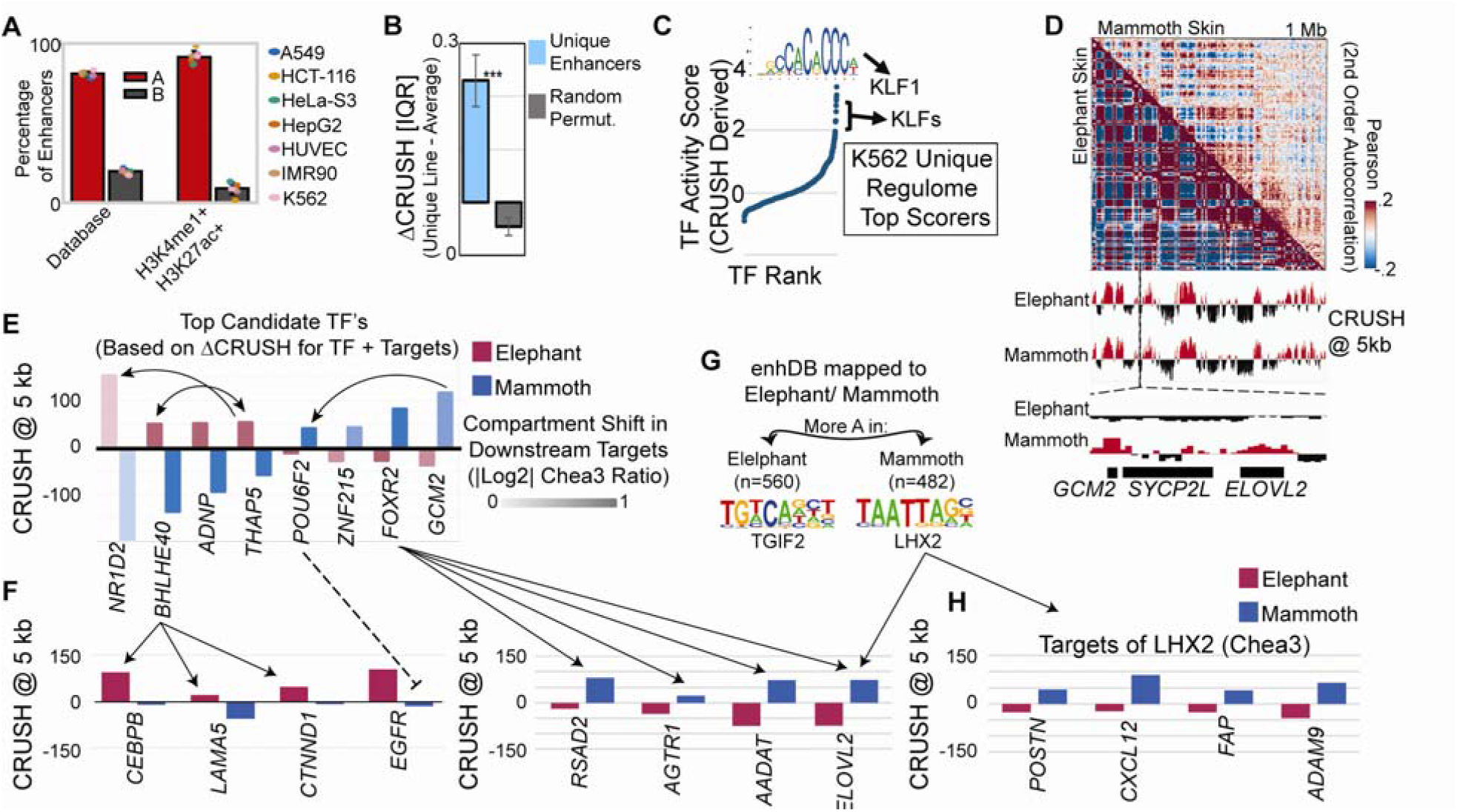
Active enhancers universally occupy the A compartment, and the woolly mammoth encodes a cold-adaptation regulome in its compartment landscape. **(A)** Enhancers from the enhDB (left) and those further filtered by the presence of H3K4me1 and H3K27ac (right) and the percentage in each cell line that localize to A (red) or B (grey) compartments binned at 1 kb. **(B)** Difference in IQR normalized CRUSH values at enhancers unique to each cell line v.s. the average CRUSH value across cell lines. *** p<.001 based on Monte Carlo permutations (grey). Error bars represent standard deviation. **(C)** TF ranks in K562 cells derived from the activity score calculated using the uniqueness of CRUSH scores at JASPAR motifs as well as at known gene targets determined by Chea3. **(D)** Example of the compartment pattern in mammoth (top right) v.s. elephant (bottom left) skin, and the resultant compartment calls at 5 kb resolution. Bottom: View of *GCM2* and *ELOVL2* which show different compartment patterns in elephant v.s. mammoth. **(E)** Top candidate TFs with different compartment patterns in elephant (red) v.s. mammoth (blue) and the log2 Chea3 scores (intensity) based on the differences in compartments on downstream targets. Arrows indicate known regulatory connections. **(F)** Chea3 identified targets of BHLHE40 and the repressor POU6F2 (left), as well as FOXR2 (right), and the corresponding compartment signal in elephant (red) v.s. mammoth (blue). **(G)** Top Homer motif in human enhancers that were mapped to the mammoth/elephant genome and identified as more active in elephant (left) or more active in mammoth (right). **(H)** Chea3 identified targets of LHX2, and the corresponding compartment signal in elephant (red) v.s. mammoth (blue).

This framework, using compartment state to infer the active regulatory landscape, is most powerfully illustrated in a context where no other epigenomic data exists. We previously annotated compartments at 50 kb in a Hi-C map from a 52,000-year-old woolly mammoth skin sample (*29*). We now resolve these compartments to 5 kb, a ten-fold improvement, and the resulting compartment signal correlates with ancient RNA recovered from a separate mammoth muscle dataset (*30*), validating biological coherence despite the degraded source material (Fig. 4D, S9C-E). Mammoth skin compartment profiles were most similar to elephant skin among all tissues examined, confirming tissue-of-origin fidelity (Fig. S9F).

To identify the regulatory programs that distinguish mammoth from elephant skin, we took two complementary approaches. First, we identified transcription factors whose encoding genes showed differential compartment state between species, with secondary scoring of these by the compartment differences of their Chea3-predicted targets (*31*) (Fig. 4E, Table S9). Elephant-specific A compartment TFs included *THAP5* and *BHLHE40*, which Chea3 states regulate each other, and whose targets, *CEBP, LAMA5*, and *CTNND1*, are associated with UV response, keratinocyte differentiation, and epidermal barrier function (*32*–*34*) appropriate for a warmer climate (Fig. 4F, left). Mammoth-specific A compartment TFs included *GCM2* and its target *POU6F2*, which can act as a transcriptional repressor (*35*) and whose motifs appear in the *EGFR* promoter and therefore may account for *EGFR*’s B-compartment localization in mammoth skin (*29*) (Fig. 4E-F, S9G). *FOXR2* was also mammoth-specific, with downstream targets *RSAD2, AGTR1, AADAT*, and *ELOVL2* having established roles in thermogenesis, vasoconstriction, melatonin signaling, and homeoviscous adaptation to cold (*36*–*39*), all in the A compartment specifically in mammoth (Fig. 4F, right). These targets are consistent with genes under positive selection in woolly mammoth identified by comparative genomic analysis (*30, 40*).

As a second, complementary approach that captures enhancer usage independent of TF gene expression, we mapped human enhancers onto the mammoth and elephant genomes and identified those with species-specific A compartment signatures. The top motif at elephant-specific enhancers was *TGIF2*, associated with keratinocyte turnover and UV response (*41*) (Fig. 4G, Table S10). The top motif at mammoth-specific enhancers was *LHX2*, a regulator of hair follicle development (Fig. 4G-H, Table S11) (*42*). Together, both regulatory inference approaches converge on the finding that the mammoth genome encodes a cold-adaptation regulome, encompassing insulation, thermogenesis, and hair follicle programs, that is written into its compartment landscape and absent from its closest living relative (see Supplementary Text).

## Conclusions

The results presented here support a revised model of chromatin compartmentalization in which the fundamental unit is not the megabase domain but the individual regulatory element. At kilobase resolution, A compartment identity precisely demarcates active enhancers and paused promoters, features that are obscured or misassigned by conventional eigenvector-based analysis (Supplemental Text). This fine-scale compartment landscape is not static as it shifts between cell types, changes during differentiation and disease progression, and is directly coupled to the transcriptional machinery through RNAPII occupancy. That these dynamics are recoverable from individual cells, without aggregation, establishes compartments as a cellular-identity that is useful for a single-cell readable dimension of chromatin state.

The breadth of biological contexts in which kilobase-scale compartments prove informative from ENCODE cell lines, (*43*) to breast cancer subtypes (*10*), and from single-cells (*24*) to a 52,000-year-old mammoth (*29*), reflects a general principle that the regulatory genome is spatially organized at the scale at which it operates. Compartment analysis has long been treated as a coarse annotation step, useful for describing global chromatin architecture but too imprecise to illuminate gene-level regulation (*2, 6*). The findings here show that this limitation was methodological rather than fundamental. With fine-scale compartment analysis now accessible in standard-depth datasets using CRUSH, nuclear organization can be examined as a routine, mechanistically interpretable dimension of genome function, one that captures the regulome of a cell, the identity of a tissue, and, as the mammoth data suggest, the adaptive logic of a species.

## Acknowledgments

We thank all the authors who created the myriad datasets used in this manuscript. We also thank individuals for helpful discussions during the creation of CRUSH and its application to uncover fundamental biological principles. Particularly helpful were discussions with the Phanstiel lab at UNC during the beginning development period, as well as Dr. Erez Aiden and Dr. Olga Dudchenko when refining the algorithm to work with ancient genomes.

## Author contributions

Conceptualization: AK, MJR

Methodology: AK

Software: AK, DU

Investigation: AK, BN, HLH, TER

Funding acquisition: MJR

Supervision: CTC, MJR

Writing – original draft: AK, MJR

Writing – review & editing: AK, CTC, MJR

## Data, code, and materials availability

Data is available as supplemental materials, with code available on the CRUSH page at github.com/JRowleyLab/CRUSH. Accessions can be found in Tables S1, S2, S4. Time and memory requirements of CRUSH are found in Table S12.

