## Supplementary material for "Kilobase-scale compartments enabled by CRUSH reveal regulatory programs across cell types, single-cells, and ancient mammoths": Supllementary Materials

Supplementary Materials

Materials and Methods

**CRUSH: Compartment Refinement for Ultraprecise Stratification of Hi-C**

CRUSH infers high-resolution chromatin compartment scores from Hi-C contact matrices using a hierarchical, multi-resolution refinement framework. Rather than relying on eigenvector decomposition of the full contact matrix, which captures global chromosomal variance and suppresses local compartment structure, CRUSH directly measures each genomic bin's interaction affinity for A-type versus B-type nuclear environments, iteratively refining these measurements from coarse to fine resolution. The algorithm proceeds in two phases: initialization of compartment reference environments at coarse resolution, followed by refinement across a resolution ladder down to 1 kb.

***Initialization.*** For each chromosome, CRUSH constructs a stabilized observed-over-expected (O/E) interaction matrix by computing the genome-wide mean contact frequency at each inter-bin distance d:

E[d] = (1/|Pd|) Σ{(i,j)∈Pd} Cij

where C is the n × n raw contact matrix and Pd is the set of bin pairs separated by d bins. The O/E field is then:

Mij = (Cij + 1) / (E[|i−j|] + 1)

Pseudocounts of 1 on both sides of the equation stabilize the ratio at sparsely sampled distances. The Pearson correlation matrix K over O/E row profiles is computed and decomposed into its top 10 principal components. To identify the component that best captures the A/B compartment axis, which need not be the first eigenvector, each component is sign-oriented so that positive values correspond to gene-enriched bins, and the component v* is selected that maximizes:

v* = arg max_v [corr(v, ρ) − corr(v, γ)]

where ρ is binned gene-body density and γ is GC-content density. This criterion selects the component most correlated with gene density and least correlated with GC content, robustly identifying the A compartment axis even when chromosome-arm organization or other sources of variance dominate the leading eigenvector. Initial A- and B-type reference environments are defined as bins with positive and negative v* scores, respectively.

***Refinement*.** CRUSH refines compartment assignments across all resolutions within the binned dataset stored as .hic (*44*) or .mcool (*45*), typically comprised of {2.5 Mb, 1 Mb, 500 kb, 250 kb, 100 kb, 50 kb, 25 kb, 10 kb, 5 kb, 1 kb}, propagating reference environments top-down from each coarser level to constrain inference at the next finer scale. At each resolution, the contact matrix is rebinned and a capped, row-wise z-scored interaction field is computed:

Zij = (Fij − μi) / (σi + ε)

where Fij = min((Cij + 1)/(Ek[|i−j|] + 1), T) is the capped O/E field (default cap T = 10), μi and σ_i are the row mean and standard deviation, and ε = 10⁻⁸ prevents division by zero. Row-wise standardization decouples compartment-driven co-segregation from bin-specific coverage artifacts arising from mappability or GC bias.

***GI scoring.*** The core compartment metric is the Genomic Interaction (GI) score, which quantifies each bin's interaction affinity for A- versus B-type environments relative to the current reference sets A{k−1} and B{k−1}:

GIi = [nA · S̄A(i) − nB · S̄B(i)] / nAB

where SA(i) = Σ{j∈A} Zij, SB(i) = Σ{j∈B} Zij, and nAB = |A{k−1}| + |B{k−1}|. Normalization by nAB ensures comparability of scores across chromosomes and resolution levels. Statistical confidence for each assignment is evaluated by a two-sample Welch t-test comparing bin i's interactions with A- versus B-reference bins; resulting p-values are corrected for multiple testing using the Benjamini-Hochberg procedure (FDR threshold α = 0.05 by default).

***Cross-resolution alignment.*** When progressing from resolution r{k−1} to rk, GI scores at the finer scale may carry a slowly varying regional offset relative to the broader genomic context. CRUSH removes this bias by subtracting a local rolling-window mean computed over a window spanning three coarse-bin widths in genomic coordinates:

GIi^centered = GIi^(rk) − μ̄i

where μ̄i is the mean GI score over the W = 3×(r{k−1}/r_k) fine bins centered on bin i. This centering preserves the relative compartment ordering within each local window while removing regional drift, anchoring each bin's score to its local genomic context. Updated reference environments for the next resolution are then defined as bins with significantly positive (A) or negative (B) centered GI scores. At fine resolutions below 50 kb, a running-mean smoothing step is optionally applied over a proportionally scaled genomic window to suppress sampling artifacts in sparse contact matrices before centering.

***Sparse bin handling.*** Bins with no observed contacts receive a pseudocount floor value of 0.1 in the zero file. These bins produce near-zero Z-scores and near-zero GI scores.

***Summary***. The full algorithm can be expressed compactly as recursive estimation of a continuous nuclear interaction field: CRUSH iterates the mapping {Ak, Bk} → Φ[·; Ak, Bk] → {A{k+1}, B{k+1}} across the resolutions until the finest requested scale is reached, at which point the centered GI scores and FDR q-values constitute the final compartment field.

**Hi-C data processing**

All Hi-C (*1*) and Micro-C (*46*) datasets were either obtained as .mcool or .hic files. Those that were not binned at 1 kb resolution were reprocessed using HiC-Pro v2.7.9 (*47*) and mapped to the appropriate reference genome: hg19 for all human cell line datasets (LCL, H1, HFFc6, breast cancer lines, and all ENCODE cell lines); GRCg6a for G. gallus erythroblasts; mm10 for mouse retina and MOE single-cell Hi-C.

**Hi-C subsampling**

Subsampled contact maps were generated by uniform random selection of read pairs from the full 20.3 billion contact LCL dataset (*3*). Subsampling scripts are available at <https://github.com/JRowleyLab/HiCSampler>.

**Eigenvector-based compartment analysis**

Eigenvector-based compartments were called using juicer and POSSUMM (<https://github.com/aidenlab/EigenVector>), which performs PCA on sparse Hi-C matrices (*3*). For all cross-method comparisons, eigenvector and CRUSH compartment calls were generated from identical input matrices at matched resolutions.

**MSD and similarity matrix calculation**

Pairwise compartment similarity between samples was quantified using the mean squared difference (MSD) of CRUSH compartment scores at all bins with valid calls in both samples. For cross-resolution comparisons, scores were resampled to a common bin size prior to MSD calculation. Pairwise similarity between mammoth skin and elephant tissue compartment profiles (skin, liver, ovary, PBMC, and brain) was computed using a similarity metric (the inverse square root of the mean squared difference [1/√MSD]) of CRUSH compartment scores at matched genomic bins.

When comparing, we place CRUSH and the eigenvector on similar scales by first normalizing each by the interquartile range (IQR). This ensures that differences are evaluated proportionally to each individual track’s range of values.

**RNA-seq processing**

RNA-seq reads were aligned to reference genomes corresponding to the respective Hi-C maps using STAR (*48*). Aligned reads were converted to bigWig format using SAMtools (*49*) and deepTools (*50*)for downstream visualization and quantification.

CRUSH compartment scores were correlated with the published ancient RNA-seq signal from mammoth muscle tissue (*30*). RNA-seq reads from that study were mapped to the Hi-C mammoth genome assembly described above using STAR, and gene-level expression values were computed as TPM.

**ChIP-seq processing**

Most signal bigwig tracks were downloaded directly from ENCODE (*43*). For datasets not available through ENCODE, reads were aligned to the appropriate reference genome using Bowtie2 (*51*) with a minimum mapping quality of MAPQ > 20. Peaks were called against matched input controls using MACS2 (p < 1×10⁻⁵) (*52*). Signal tracks were generated using deepTools bamCoverage (*50*).

**Compartment intersection visualization**

Intersections of compartment assignments across ENCODE cell lines were visualized as UpSet plots computed from binary A/B assignments across all cell-line combinations. Regional compartment discrepancies between CRUSH and eigenvector were visualized using HiCrayon (*53*), with H3K27ac signal used to assess concordance with active chromatin state.

**Chromatin state annotation**

ChromHMM (*18*) segmentation files were obtained from the Roadmap Epigenomics Project (*54*). Chromatin state annotations were intersected with genomic bins at each resolution using BEDTools (*55*), with each bin assigned the state with the greatest overlap. Bins were classified as stable A, stable B, or variable based on compartment consistency across all seven ENCODE cell lines. Enrichment of each ChromHMM state in variable versus stable bins was computed as log₂(f_variable / f_stable), where f denotes the fraction of bins in each category carrying a given state. Negative values indicate depletion in variable regions; positive values indicate enrichment.

**Logistic regression analysis**

To quantify the predictive power of individual chromatin features for compartment identity, logistic regression models were trained independently at 100 kb, 40 kb, and 1 kb resolution using binary compartment status (A = 1, B = 0) as the response variable. Predictor variables included BRU-seq signal, H3K27ac, H3K4me1, H3K27me3, and a combined multi-feature model. All feature values were z-score normalized prior to modeling. Model performance was assessed by 5-fold cross-validation AUC-ROC. Signal tracks used for regression were obtained from the Avocado (*56*).

**BRU-seq and RNAPII pausing analysis**

BRU-seq signal (*43*) profiles were analyzed for genes longer than 5 kb. Paused genes were defined as those with high nascent transcription signal at the TSS and low signal across the gene body; elongating genes showed distributed signal across the full gene body. Average BRU-seq and CRUSH compartment score profiles were computed across gene bodies from -1 gene length (GL) before the TSS to +1 GL after the TTS. All signals were IQR (Inter-quartile range)-normalized to enable comparison across cell lines. (IQR-normalization entails subtracting the median and dividing by the interquartile range across all bins with valid scores within each sample.) RNAPII occupancy and compartment profiles in DLD1 RNAPII-AID cells (*19*) were analyzed under control (-auxin) and depletion (+auxin) conditions using RNAPII ChIP-seq generated with an anti-GFP antibody against the endogenously tagged RPB1.

**Single-cell compartment analysis**

Single-cell DIP-C contact maps from mouse retina, MOE, PBMCs, and GM12878 cells (*23*, *24*) were processed with CRUSH at 500 kb, 100 kb, 50 kb, and 25 kb resolution. Reference A/B environments for CRUSH initialization were derived from pseudo-bulk Hi-C maps of the corresponding cell types at 100 kb; these bulk-derived states were used solely for first round initialization and were excluded from all downstream single-cell analyses. Per-cell compartment scores across all genomic bins were assembled into a cell × bin matrix. Features were z-score normalized and UMAP was applied independently at each resolution (cosine metric, 15 neighbors, minimum distance 0.01, random seed 42). Cell-type-specific gene-compartment signatures were defined using the top 150 genes with the largest differential compartment scores between populations. Classifier performance was evaluated by AUC-ROC.

Single-cell clustering quality was assessed using the neighborhood purity index at k = 15, defined as the fraction of the k nearest neighbors in UMAP space belonging to the same cell-type label as the query cell, averaged across all cells:

NPI = (1/|C|) Σ_{c∈C} (1/k) Σ_{j∈kNN(c)} 𝟙[label(j) = label(c)]

Values range from 0 (completely mixed populations) to 1 (perfect separation). NPI was computed independently at each resolution to quantify the resolution-dependence of cell-type segregation.

**TCGA expression analysis**

Breast cancer gene expression data across subtypes (Normal, Luminal A, Luminal B, HER2+, and TNBC) were obtained via the UALCAN portal (*12*), which provides access to TCGA RNA-seq data. Expression levels for genes of interest were compared across subtypes and integrated with CRUSH and eigenvector compartment scores to assess concordance between compartment state and patient-level transcriptional activity.

**Transcription factor activity scoring in mammoth and elephant skin**

To identify transcription factors with differential regulatory activity between mammoth and elephant skin (*29*), we used two complementary approaches. First, TF-encoding genes were ranked by the difference in their CRUSH compartment scores between species (|ΔCRUSH| ≥ 50), weighted by a ChEA3 (*31*) derived regulon score reflecting the compartment score differences of their predicted downstream targets. ChEA3 was run with default parameters in integrated mean rank mode, which aggregates evidence across co-expression, ChIP-seq, and reporter assay libraries, querying gene sets defined by differential A compartment state between species. The composite TF score thus reflects both the compartment state of the TF gene itself and the collective compartment state of its regulon, prioritizing factors whose entire regulatory program shifts between species.

As a complementary enhancer-centric approach, 1 kb sequences corresponding to human enhancers from the Enhancer Database were mapped onto the mammoth and elephant genome assemblies us minimap2 (*57*). Putative enhancers with a CRUSH compartment score difference of ≥ 50 between species were designated as mammoth- or elephant-specific. Enriched TF binding motifs at species-specific enhancers were identified using HOMER findMotifsGenome.pl (*58*).

**Transcription factor activity scoring in ENCODE cell lines**

Cell-line-specific enhancers were defined using annotations from the Enhancer Database (*27*) and further refined by overlap with cell-line-matched H3K27ac and H3K4me1 ChIP-seq peaks. TF binding motifs at these enhancers were obtained from the JASPAR database (*59*). For each TF motif and each cell line, a compartment-enhancer activity score was computed as the mean difference between the IQR-normalized CRUSH score at motif-containing enhancers in that cell line and the average IQR-normalized CRUSH score at the same enhancers across all seven ENCODE lines, quantifying how distinctively active those enhancers are in each cellular context.

In parallel, genes with an IQR-normalized CRUSH score ≥ 50 above the mean of the other six lines were defined as uniquely compartmentalized in that cell line and submitted to ChEA3 (*31*) to rank TFs by overrepresentation of uniquely A compartment targets. Final TF activity scores were computed as the mean of the motif-based compartment score rank and the ChEA3 target-based rank. These composite scores were normalized within each cell line by computing the −log₂ ratio relative to the mean score across all seven lines, yielding a cell-line-specific TF activity metric that reflects both enhancer usage and downstream target compartmentalization.

**Gene Ontology analysis**

GO Biological Process enrichment analysis was performed using EnrichR (*60*) on genes overlapping bins with compartment differences, with BH-corrected p-values. Significance of top GO terms identified by CRUSH versus eigenvector was compared using a two-sided Wilcoxon signed-rank test.

Supplementary Text

Eigenvector decomposition versus CRUSH, a technical comparison

Eigenvector-based compartment calling relies on principal component analysis (PCA) on the Pearson correlation matrix of observed-over-expected (O/E) Hi-C contact frequencies (*1*, *6*). The first eigenvector (PC1) is then used as a proxy for the A/B compartment axis, with sign orientation determined by manual inspection of each chromosome or post-calculation correlation with gene density or GC content (*6*). This approach has two fundamental limitations that compound at high resolution. First, PCA maximizes global variance across the chromosome, not the variance attributable to the A/B compartment axis specifically. When other sources of genomic variance, most notably chromosome-arm-level contact frequency gradients, or large-scale copy number variation in cancer genomes, account for more variance than compartment identity, PC1 captures those signals instead (*6*). The compartment pattern, which in such cases may be clearly visible as a checkerboard in the raw contact map, is displaced to a higher component that is never reported. Second, because eigenvector decomposition is performed on the full-resolution contact matrix at once, signal-to-noise scales unfavorably with resolution because the fraction of the Pearson correlation matrix entries supported by observed contacts drops as bin size decreases, so the eigenvector becomes increasingly dominated by sampling noise below approximately 25 kb (*3*). Achieving stable 1 kb eigenvector calls requires contact depths on the order of billions of intrachromosomal pairs (*3*), placing fine-scale compartment analysis out of reach for the vast majority of published datasets.

CRUSH addresses both limitations through a fundamentally different inferential strategy. Rather than decomposing the full contact matrix globally, CRUSH measures each bin’s local interaction affinity for A-type versus B-type reference environments, the Genomic Interaction (GI) score, and refines these measurements iteratively across a resolution ladder. Only the very first initialization step applies PCA to the coarsest-resolution O/E Pearson matrix. but also uses a component-selection criterion that explicitly maximizes correlation with gene density while minimizing correlation with GC content, rather than defaulting to PC1. This component-selection step is what allows CRUSH to recover the correct axis in genomes where eigenvector misassignment occurs; therefore, by scoring all top components against biological priors and selecting the best-aligned one, CRUSH is robust to the variance-dominance of arm-level or copy-number signals that mislead the eigenvector. At subsequent refinement steps, reference environments are propagated top-down from coarser to finer resolution, as well as bottom-up from finer to coarser resolutions, so that each fine-scale GI score is computed against reference sets that are already well-characterized from higher-depth, lower-resolution bins as well as from lower-depth, high-resolution bins. The result is that CRUSH achieves stable 1 kb compartment calls at contact depths two to three orders of magnitude below what the eigenvector requires to approach comparable precision, as demonstrated by the subsampling experiments in Fig. 1A and S1B-D. Although outside the scope of this work, in the future, we imagine differential compartment callers like dcHiC (*61*), which currently rely on the eigenvector, may be improved if retooled to use IQR normalized CRUSH, as the low sequencing threshold and inter-replicate consistency makes utilizing replicates in the statistical determination of differences more feasible.

A further difference between the two methods is how they handle cross-methodology bias. Micro-C and Hi-C use different crosslinking chemistries and fragment size distributions that alter the short-range contact frequency profile and shift the effective O/E background at distinct distances (*3*, *7*, *46*, *62*, *63*). Because the eigenvector operates on global Pearson correlations computed from the full interaction distance range, these protocol-dependent short-range effects propagate into the compartment calls, producing systematic offsets between Hi-C and Micro-C maps from the same cell line that can exceed the biological differences between cell types (Fig. 1B-C, S1E). CRUSH’s row-wise z-scoring of the O/E interaction field prior to GI scoring decouples compartment-driven co-segregation signals from bin-specific coverage artifacts, including those arising from protocol-dependent short-range enrichment, because normalization is computed independently per bin rather than across the full matrix. This is why CRUSH recovers consistent compartment calls across Hi-C and Micro-C while the eigenvector does not.

Compartment dynamics during erythroid maturation in *G. gallus*

In maps of *G. gallus* erythroblasts (*9*), the eigenvector consistently fails to report A/B compartments, instead capturing chromosome-arm-level contact frequency gradients as its leading principal component (Fig. S2A). This failure is not a feature of the underlying chromatin organization, the checkerboard compartment pattern characteristic of A/B segregation is clearly visible in the contact map at 50 kb resolution, but reflects the eigenvector’s global variance maximization, encountering a genome in which non-compartment contact enrichment dominates the Pearson correlation structure. Because CRUSH performs the first initialization (and only the first initialization) by selecting the component most aligned with gene density rather than defaulting to PC1, and then iteratively corrects the initialization by direct measurement of each row’s interaction preferences, it recovers correct compartment assignments at both 50 kb and 1 kb resolution, with compartment calls that show substantially improved concordance with RNA-seq expression levels compared to the eigenvector at matched resolution (Fig. S2C).

Applying CRUSH across fibroblast, embryonic RBC (eRBC), erythroblast, and adult RBC (aRBC) stages reveals a structured compartment transition trajectory during erythroid maturation (Fig. S2B, F). The number of compartment switches increases progressively from the fibroblast-to-erythroblast transition through terminal erythroid differentiation, with the erythroblast-to-eRBC and eRBC-to-aRBC transitions each showing a distinct set of loci undergoing A-to-B or B-to-A conversion. Loci switching from A to B during terminal differentiation, such as *KANK1* and *GNAI1* (Fig. S2G), are transcriptionally downregulated in the RNA-seq data, consistent with compartment-driven silencing of genes not required in the mature erythrocyte. Enrichment analysis of loci undergoing compartment transitions across the erythroid trajectory recovers pathways relevant to each stage, RAP1 signaling, proteoglycan remodeling, and cell cycle withdrawal in early erythropoiesis; and cellular senescence programs in the terminal eRBC-to-aRBC step (Fig. S2E, H). These stage-specific compartment transitions were not recoverable by the eigenvector at any resolution tested, underscoring that the biological signal captured here depends directly on the correction of compartment misassignment. The systematic nature of this failure, not random noise but a structured misdirection toward a non-compartment variance axis, reinforces the importance of component selection as a prior step in compartment analysis of non-standard or highly structured genomes.

RNAPII pausing and the determinants of sub-genic compartment identity

The RNAPII depletion experiment in DLD1 RNAPII-AID cells (*19*) demonstrates that polymerase occupancy is causally required for the TSS-proximal A compartment spike observed at paused genes. However, the incomplete conversion of paused gene bodies to the B compartment following auxin-induced RNAPII degradation (Fig. S4D) indicates that polymerase occupancy is only one of multiple determinants of local compartment identity. Several chromatin features that are independent of active transcription are likely contributors. H3K4me3-marked promoters retain some degree of A compartment affinity even in the absence of active elongation, consistent with prior observations that promoter-associated histone modifications persist through transcriptional inhibition and may maintain local chromatin accessibility independent of polymerase activity. The observation that nascent transcription (BRU-seq) is the strongest single predictor of 1 kb compartment identity across cell lines (Fig. 2C-D, S5C) does not preclude these additional determinants; rather, it reflects that transcription integrates and correlates with many of the same chromatin features that independently contribute to compartment state. Disentangling the causal relationship among polymerase occupancy and other 1D active marks as determinants of sub-genic compartment identity will require combinatorial perturbation experiments beyond the scope of the current study.

Reference environment initialization in single-cell compartment analysis

CRUSH requires initial A- and B-type reference environments to anchor GI score computation at the coarsest resolution. In bulk Hi-C analysis, these reference sets are derived from the coarse-resolution Pearson PCA of the dataset itself. In single-cell Hi-C maps, contact coverage per cell is typically two to four orders of magnitude lower than bulk experiments, making reliable PCA-based initialization from individual cell maps intractable at any resolution. To circumvent this, we derived initial A/B reference environments from pseudo-bulk Hi-C maps of the corresponding cell types at 100 kb resolution. These pseudo-bulk-derived reference states were used solely to define the first (and only the first) initialization state and were explicitly excluded from all downstream single-cell analyses so that per-cell GI scores were computed exclusively from the individual cell’s own contact matrix, and the pseudo-bulk reference was not consulted after the first initialization.

This design choice has an important methodological consequence; because the reference environments are shared across all cells in a dataset, they cannot themselves drive cell-type separation in the resulting UMAP. Any clustering that emerges from per-cell CRUSH profiles must reflect genuine variation in individual cells’ interaction affinities for those reference environments, not artifacts of cell-type-specific initialization. This approach is analogous in principle to the use of a shared reference genome for single-cell RNA-seq quantification, as the reference defines the coordinate system but does not determine which cells express which genes. The per-cell compartment scores reported here reflect each individual cell’s nuclear organization within that shared coordinate framework.

The mammoth cold-adaptation regulome, transcription factor networks, and biological context

The two complementary regulatory inference approaches applied to mammoth and elephant skin, TF gene compartment scoring weighted by ChEA3 regulon analysis (*31*), and enhancer-centric motif enrichment at species-specific A compartment enhancers, converge on overlapping but distinct sets of biological programs, which warrants extended interpretation. The TF-centric approach identifies factors whose own genes are differentially compartmentalized between species and whose predicted downstream targets show concordant compartment shifts, prioritizing regulators whose entire transcriptional program is reorganized. The enhancer-centric approach instead captures regulatory elements that are active in one species but not the other, independent of whether the TF gene itself is differentially compartmentalized and is therefore sensitive to cases where a shared TF drives species-specific programs through differential enhancer accessibility rather than differential TF expression.

The mammoth-specific FOXR2 regulon identified by the TF-centric approach includes four targets with well-characterized roles in cold physiology. *RSAD2* (viperin) has established functions in lipid droplet biogenesis and has been shown to modulate thermogenic capacity in adipocytes (*36*). AGTR1 (angiotensin II receptor type 1) mediates vasoconstriction and is a key effector of peripheral vascular tone regulation in cold exposure , consistent with the selective vasoconstriction needed to minimize heat loss in arctic environments (*37*, *64*). *ELOVL2* encodes a fatty acid elongase that extends very long-chain polyunsaturated fatty acids (VLCPUFAs), a process central to homeoviscous adaptation, the maintenance of membrane fluidity at low temperatures through VLCPUFA enrichment (*39*, *65*). The co-occurrence of these targets in the mammoth A compartment, absent from elephant, and their functional coherence around cold physiology, supports the interpretation that FOXR2 coordinates a cold-adaptation transcriptional program that is encoded in the mammoth regulatory genome.

The enhancer-centric analysis independently identifies LHX2 as the top motif at mammoth-specific A compartment enhancers. LHX2 is a homeodomain transcription factor with well-documented roles in hair follicle morphogenesis and cycling, and it is required for the establishment of the dermal papilla niche and the maintenance of the anagen (growth) phase of the hair cycle (*42*). The convergence of both regulatory inference strategies on cold-adaptation themes (thermogenesis, vasoconstriction, lipid metabolism, and hair follicle elaboration) in the mammoth, and on UV response and keratinocyte programs in the elephant, provides mutually reinforcing evidence that these compartment differences reflect genuine adaptive divergence in the regulatory genomes of the two species rather than stochastic genomic drift.

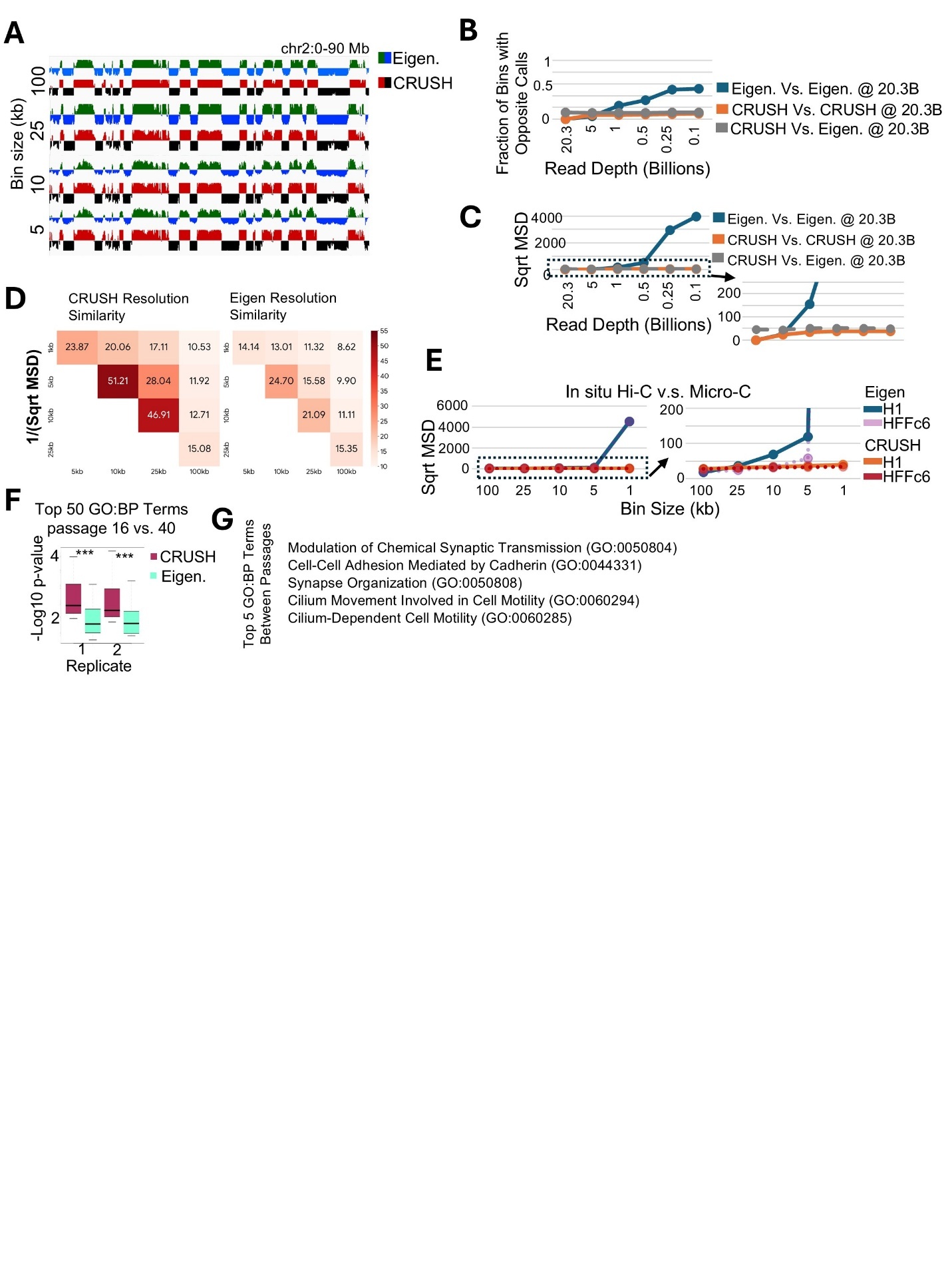

Fig. S1.

**(A)** Comparison of the eigenvector and CRUSH at 100kb, 25kb, 10kb and 5 kb resolution in LCLs with 20.3 billion intra-chromosomal Hi-C contacts. **(B)** Fraction of bins with opposite calls comparing CRUSH (orange) and the eigenvector (blue) in subsampled maps to their respective calls in the full 20.3 billion contact map. Also shown is a comparison of CRUSH at each sequencing depth to that of the eigenvector in the full 20.3 billion contact map (grey). **(C)** Sqrt Mean Squared Difference (MSD) of CRUSH (orange) and the eigenvector (blue) in subsampled maps to their respective calls in the full 20.3 billion contact map. Also shown is a comparison of CRUSH at each sequencing depth to that of the eigenvector in the full 20.3 billion contact map (grey). **(D)** 1/sqrt (MSD) similarity matrices between CRUSH and eigenvector-derived calls across resolutions. **(E)** Sqrt MSD at each resolution using the eigenvector in H1 (blue) and HFFc6 (purple) and CRUSH in H1 (orange) and HFFc6 (red) Hi-C compared to their respective calls in Micro-C. Dashed rectangle indicates that portion of the graph zoomed-in on the right. **(F)** Significance scores of the top 50 Gene Ontology Biological Process (GO:BP) terms associated with differential compartments across passages when using CRUSH vs Eigenvector. *** p<.001 Wilcoxon rank sum test. **(G)** Top 5 GO:BP terms associated with differential compartments across Wi38 cell passages as identified by CRUSH.

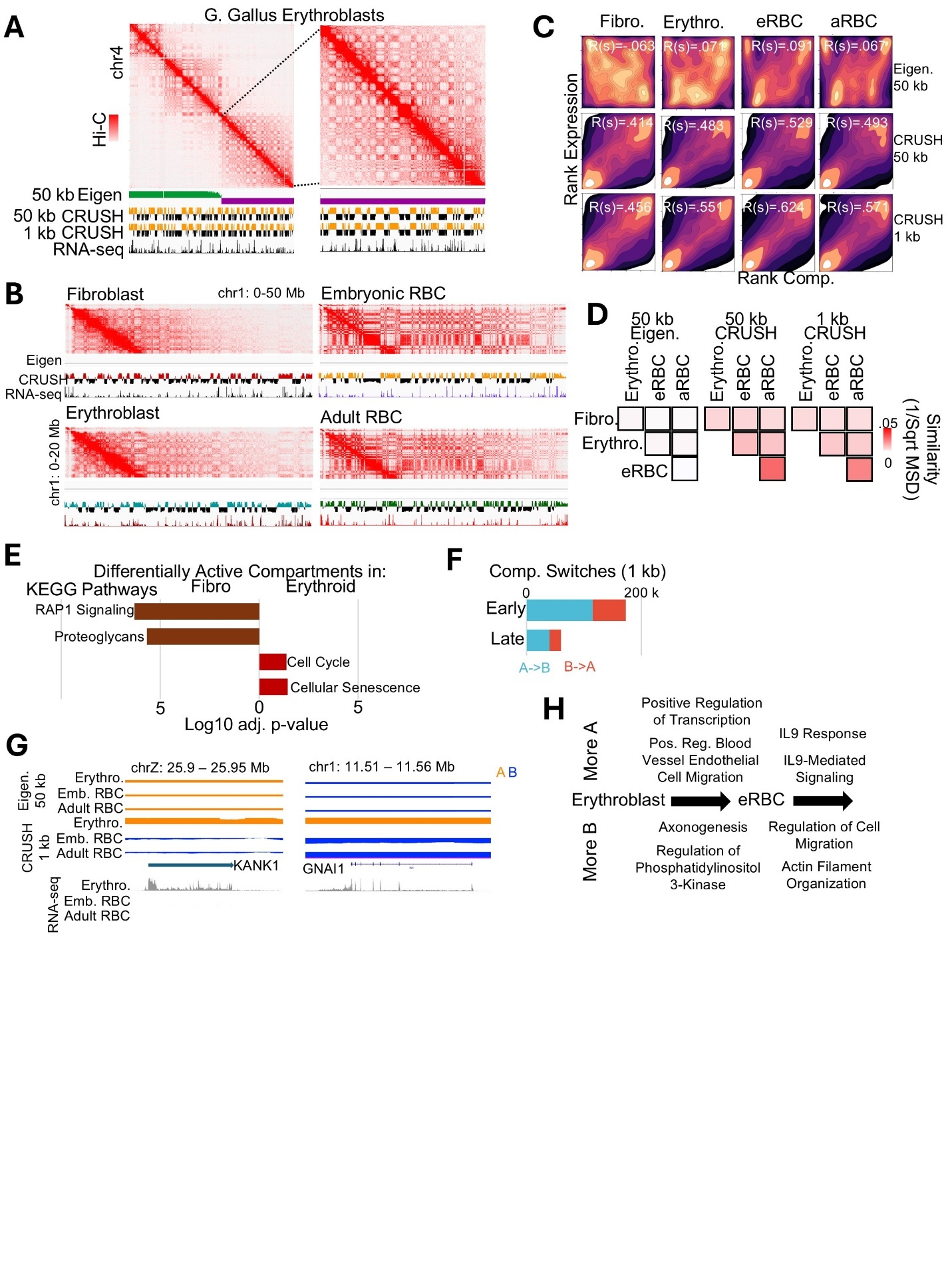

Fig. S2.

**(A)** Eigenvector at 50 kb compared to CRUSH at 50 kb and 1 kb resolution, along with RNA-seq and the Hi-C map in *G. gallus* erythroblasts, to showcase how the separation of chromosome arms can impede the eigenvector from detecting compartments. **(B)** Hi-C checkerboard and the 1 kb CRUSH compartment tracks for chr1: 0-50 Mb across Fibroblast, Embryonic RBC, Erythroblast, and Adult RBC. **(C)** Rank expression vs. rank compartment score for the eigenvector compartments at 50 kb and CRUSH at 50 kb or 1 kb across fibroblasts (Fibro.), erythroblasts (Erythro.), as well as embryonic and adult Red Blood Cells (eRBC and aRBC). Spearman R(s) values noted. **(D)** Similarity matrices (1/sqrt MSD) comparing cell types at 50 kb Eigen, 50 kb CRUSH, and 1 kb CRUSH resolution. **(E)** KEGG pathway enrichment for differentially active compartments in fibroblast vs. erythroid lineages. **(F)** Number of A to B (blue) and B to A (red) compartment switches during the transition from erythroblasts to embryonic RBCs (early) to adult RBCs (late). **(G)** *KANK1* and *GNAI1* loci as examples of differences between erythroblasts and RBCs that were missed by the eigenvector. RNA-seq signal shown to denote expression status of the gene. **(H)** Gene Ontology: Biological Processes (GO:BP) enrichment terms for the erythroblast-to-eRBC-to-aRBC compartment switch trajectory.

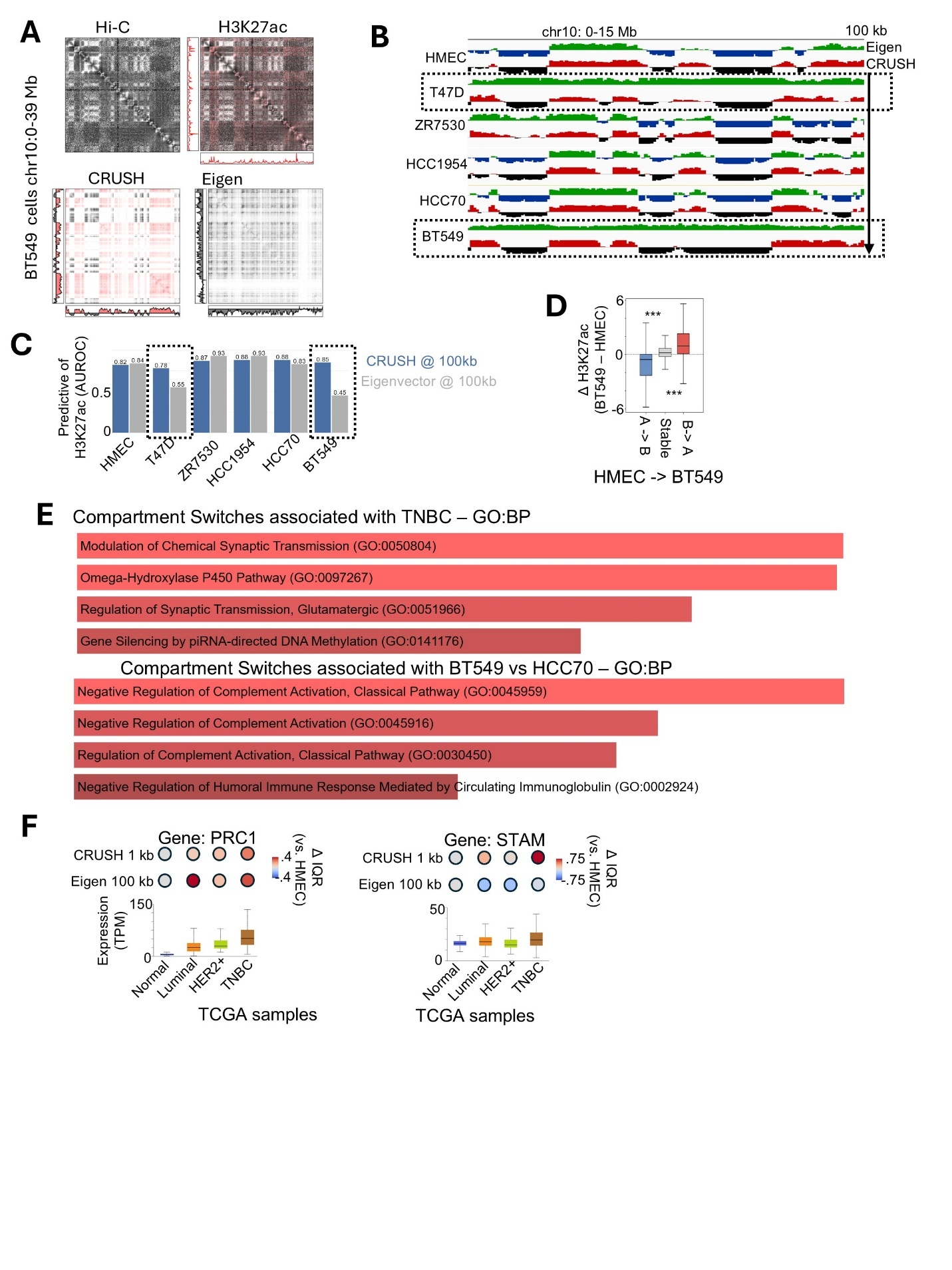

Fig. S3.

**(A)** HiCrayon visualization demonstrates that CRUSH calls match the checkerboard pattern that was missed by the eigenvector in BT549 cells. H3K27ac is shown as a representative active mark. **(B)** Compartment tracks called by the eigenvector vs. CRUSH calls at 100 kb resolution. Dashed boxes indicate cells in which the eigenvector failed to identify the compartments for this region. **(C)** The relationship between H3K27ac and compartments called by CRUSH (blue) or the eigenvector (grey). 100 kb bins were used because the eigenvector failed to produce coherent compartments at 1 kb. **(D)** Difference in H3K27ac between HMEC and BT549 at bins that switch from A to B (blue), don’t change (grey), or switch from B to A (red). **(E)** Top GO:BP terms associated with compartment differences between triple negative breast cancer (TNBC) and non-TNBC cells. Bottom: GO:BP terms associated with compartments unique to BT549 vs. HCC70. **(F)** Dot and boxplot for *PRC1* (left) and *STAM* (right) gene compartment scores (IQR normalized and relative to HMEC) in cell lines compared to expression in patients across TCGA samples categorized as Normal, Luminal, HER2+, and TNBC.

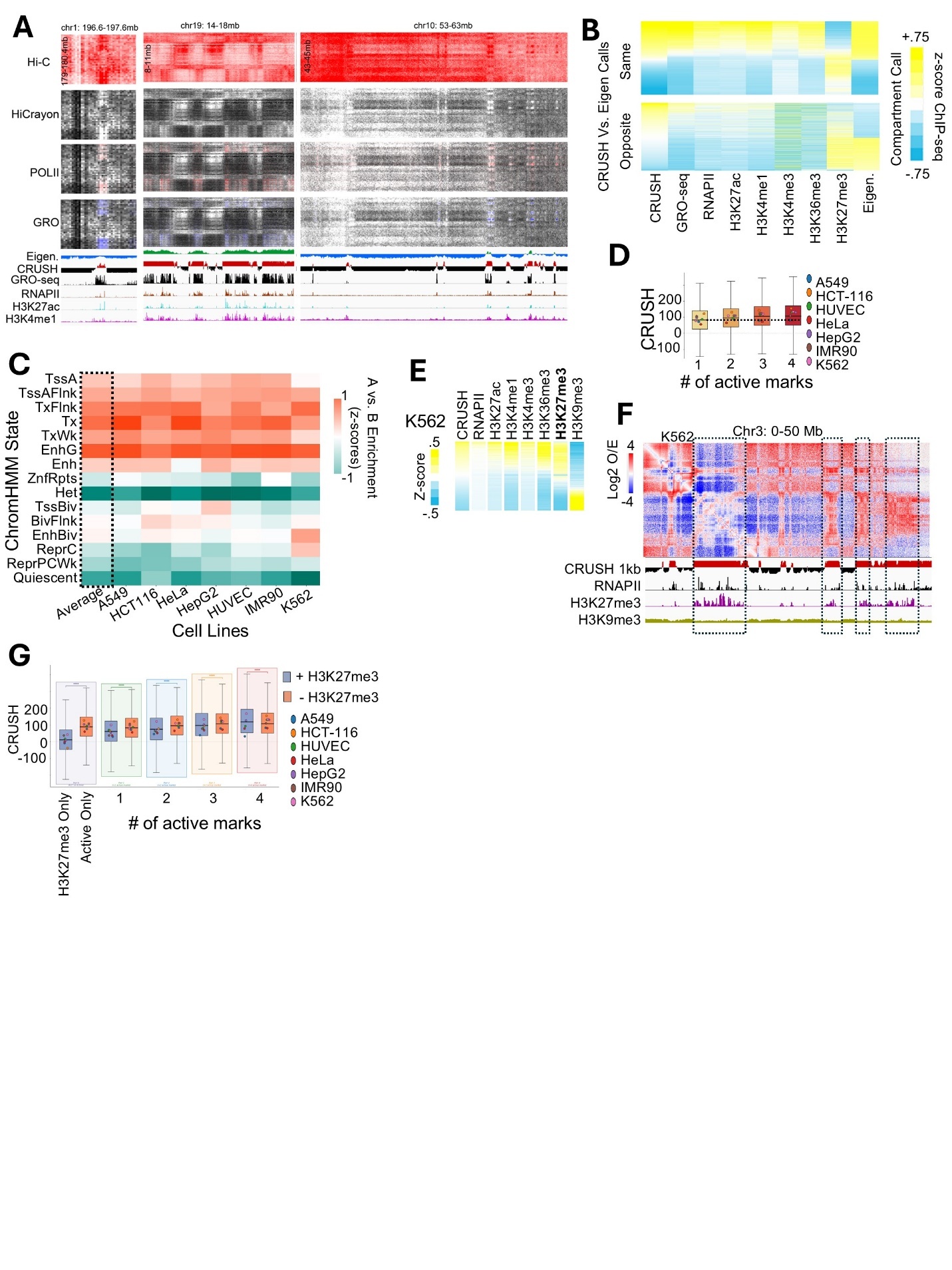

Fig. S4.

**(A)** Example of a compartment call mismatches between the eigenvector and CRUSH in the deeply sequencing LCL Hi-C map. GRO-seq, RNAPII, H3K27ac, and H3K4me1 are shown to denote 1D chromatin activity state. HiCrayon was used to color maps by the 1D features to illustrate the overlap with the compartment checkerboard. **(B)** Heatmap of bins with the same (top) or opposite (bottom) compartment call between CRUSH and the eigenvector, and chromatin features including GRO-seq, and ChIP-seq for RNAPII, H3K27ac, H3K4me1, H3K4me3, H3K36me3, and H3K27me3 (z-score normalized to allow usage of similar scales). **(C)** ChromHMM states enriched in 1 kb A or B compartments across cell lines. **(D)** CRUSH scores in 1 kb bins with different numbers of active marks. Dots represent the average of each individual cell line, while boxes represent the aggregate across cell lines. **(E)** Heatmap of 1 kb CRUSH scores vs. chromatin marks RNAPII, H3K27ac, H3K4me1, H3K4me3, H3K36me3, H3K27me3, and H3k9me3 (z-score normalized signal). H3K27me3 is bold to highlight its presence in the A compartment and the overlap with other active marks. **(F)** Example of the overlap between H3K27me3 and RNAPII, along with the association with A compartments in K562 cells (dashed boxes). Hi-C heatmap is distance-normalized (Log2 OE [observed/expected]) to highlight the checkerboard compartment pattern. **(G)** CRUSH scores in 1 kb bins with different numbers of active marks as well as with (blue) or without (orange) H3K27me3. Dots represent the average of each individual cell line, while boxes represent the aggregate across cell lines. Wilcoxon rank-sum test, *** p < 0.001.

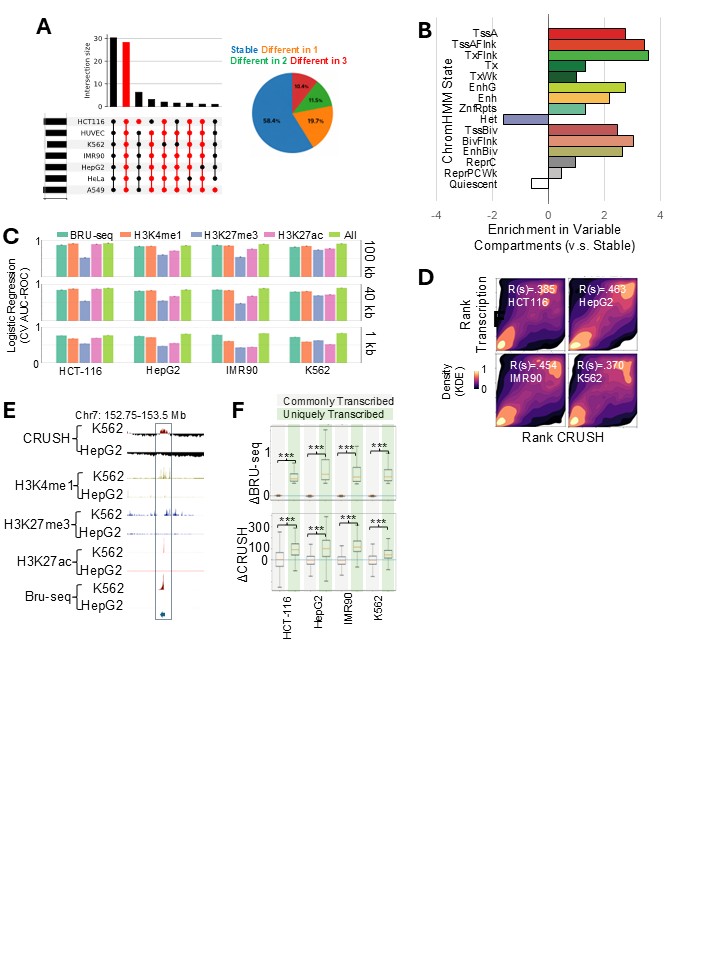

Fig. S5.

**(A)** Left: UpSet plot shows how many genomic bins share compartment calls across seven ENCODE cell lines, with bar height for intersection size and filled circles for included lines. Right: Pie chart displays the proportions of bins that are stable or differ across one, two, or three-plus cell lines. **(B)** ChromHMM state enrichment for compartments bins that were variable across seven cell lines (log2 ratio vs. stable bins). **(C)** Cross-validated logistic regression performance (AUC-ROC) for predicting compartment identity using BRU-seq, H3K4me1, H3K27me3, H3K27ac, or all marks combined at 100 kb, 40 kb, and 1 kb resolution. **(D)** Density contour plots showing the joint distribution of ranked BRU-seq transcription signal (y-axis) and ranked CRUSH compartment score (x-axis) for genes in HCT-116, HepG2, IMR90, and K562. **(E)** Example of a gene with opposite compartment association between K562 and HepG2. H3k4me1, H3K27me3, H3K27ac, and BRU-seq are shown for reference. **(F)** Differences in BRU-seq signal (ΔBRU-seq, top) and differences in CRUSH scores (ΔCRUSH, bottom) between bins transcribed in all four cell lines and those unique to the indicated line. Wilcoxon rank-sum test, *** p < 0.001.

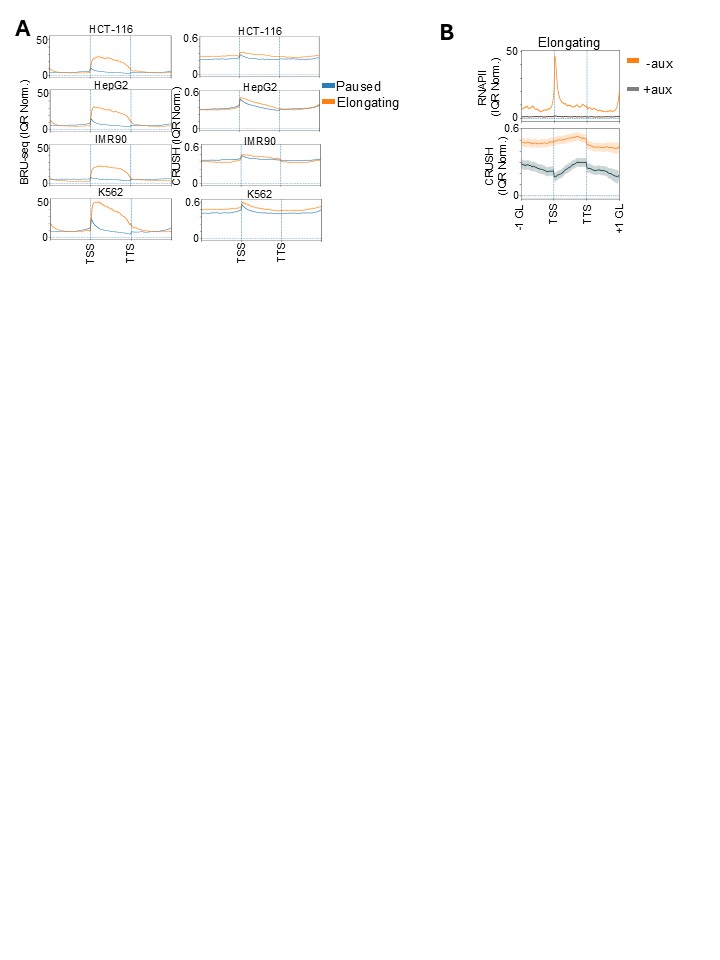

Fig. S6.

**(A)** Average BRU-seq (left) and CRUSH score (right) profiles across gene bodies for paused (blue) and elongating (orange) genes, going −1 gene length (GL) before the TSS to +1 GL after the TTS. Profiles are shown as IQR-normalized means in each cell line. Shaded bands represent standard deviation. **(B)** RNAPII occupancy (top) and CRUSH scores (bottom) in DLD1 RNAPII-AID cells in -auxin (orange) and +auxin (grey) conditions. Shaded bands represent standard deviation.

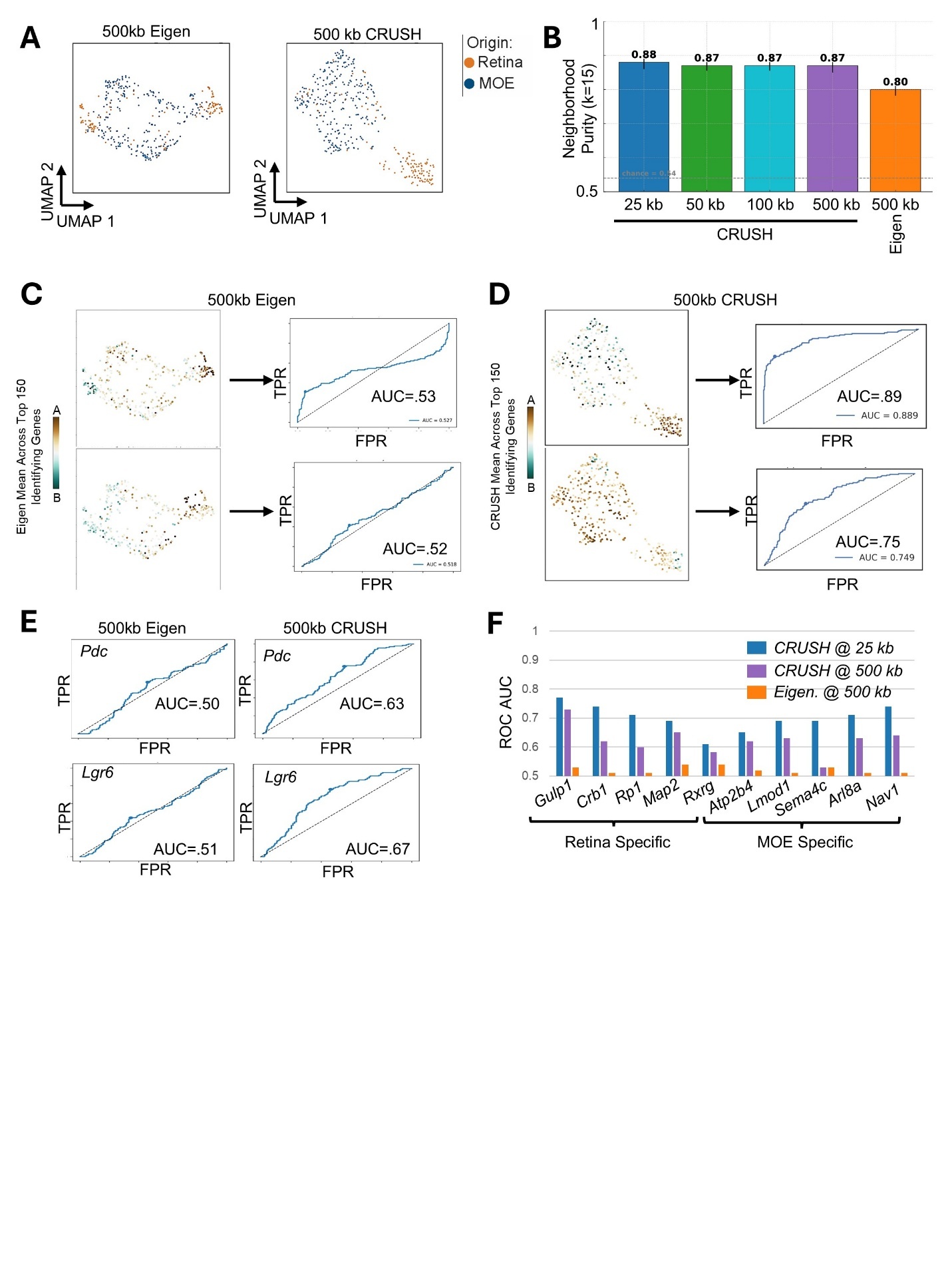

Fig. S7.

**(A)** DIP-C data comprised of retina (orange) and MOE (blue) with the UMAP based on 500 kb eigenvector (left) or 500 kb CRUSH (right). **(B)** Neighborhood purity index (k=15) for CRUSH at various resolutions alongside the eigenvector at 500 kb. Grey dashed line indicates random permutations. **(C)** 500 kb eigenvector-based compartments and the corresponding UMAP colored by top 150 genes with unique A compartment signatures in the retina (top) and MOE (bottom) alongside the Receiver Operating Characteristic (ROC) based on the True Positive Rate (TPR) and False Positive Rate (FPR). AUC = Area Under Curve. **(D)** 500 kb CRUSH-based compartments and the corresponding UMAP colored by top 150 genes with unique A compartment signatures in the retina (top) and MOE (bottom) alongside the ROC. AUC = Area Under Curve. **(E)** AUC ROC for *Pdc* (top) and *Lgr6* (bottom) using compartments from the 500 kb eigenvector (left) or 500 kb CRUSH (right). **(F)** AUC for 5 retina-specific and 5 MOE-specific gene-compartment signatures calculated by CRUSH at 25 kb (blue), CRUSH at 500 kb (purple), or the eigenvector at 500 kb (orange).

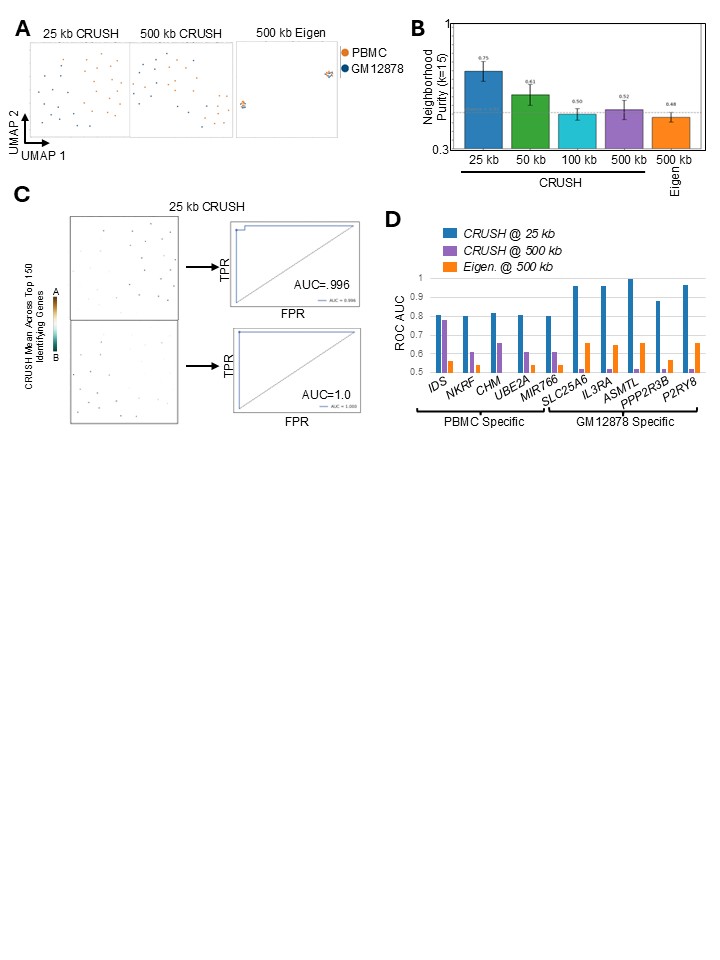

Fig. S8.

**(A)** Single-cell compartment analysis of mixed PBMC (orange) and GM12878 (blue) cells by CRUSH at 25 kb (left), CRUSH at 500 kb (middle), and eigenvector at 500 kb (right) with UMAP visualization. **(B)** Neighborhood purity index (k=15) for CRUSH at various resolutions for PBMC/GM1878 alongside the eigenvector at 500 kb. Grey line indicates value at random distribution. **(C)** Left: UMAP colored by top 150 genes with unique A compartment signatures in PBMCs (top) and GM12878 (bottom). Right: Receiver Operating Characteristic (ROC) based on the True Positive Rate (TPR) and False Positive Rate (FPR) for the top 150 compartment signatures. AUC = Area Under Curve. **(D)** AUC ROC for 5 PBMC-specific and 5 GM12878-specific gene-compartment signatures calculated by CRUSH at 25 kb (blue), CRUSH at 500 kb (purple), or the eigenvector at 500 kb (orange).

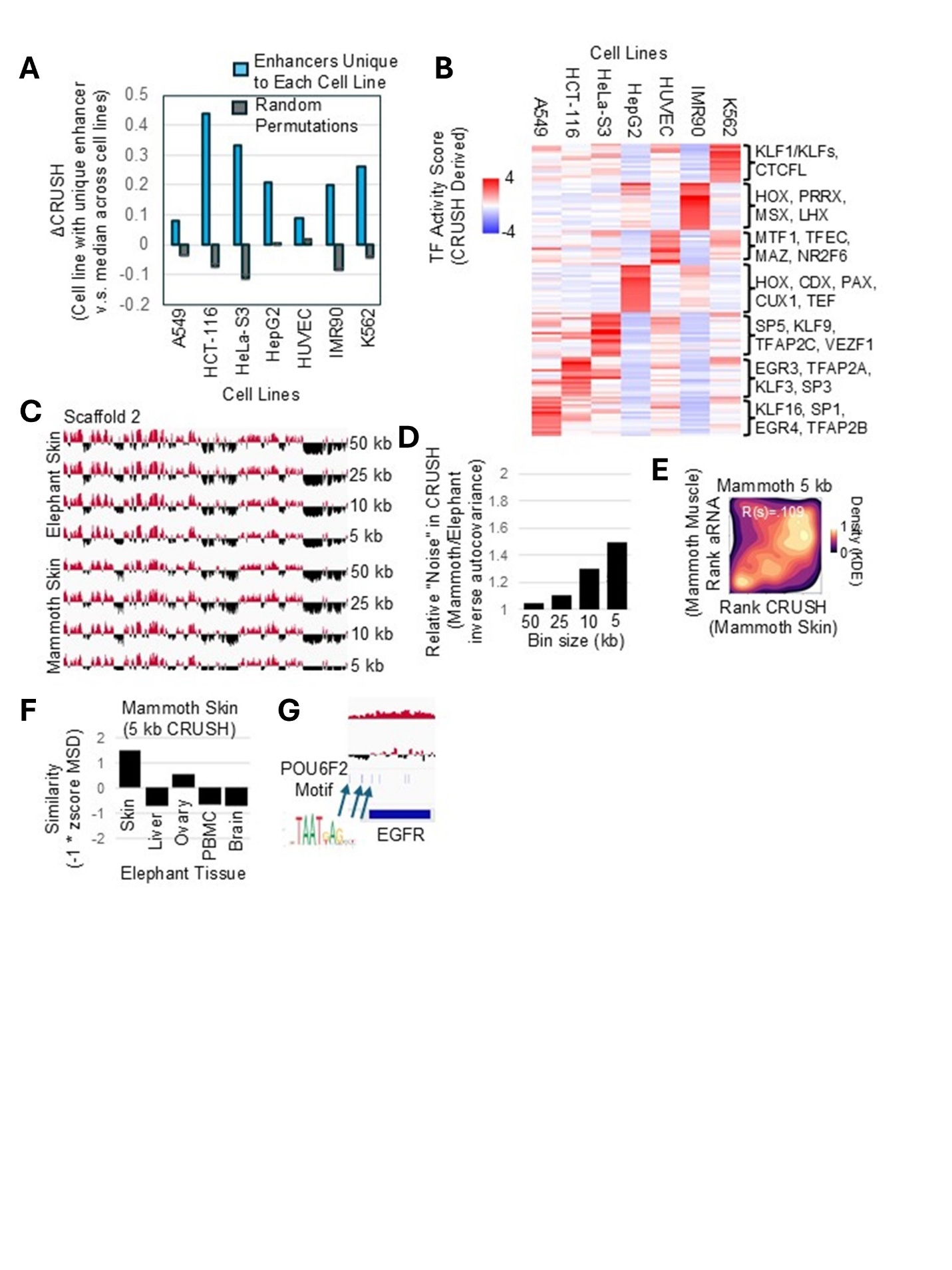

Fig. S9.

**(A)** Difference in IQR normalized CRUSH values at enhancers unique to each cell line vs. the median across cell lines. Monte Carlo 1000 permutations (grey). **(B)** TF activity score derived from CRUSH for enhancers with unique compartment signatures in each cell line. **(C)** Refinement of compartments from 50 kb to 5 kb resolution in elephant and mammoth skin Hi-C maps. **(D)** An estimate of the relative “noise” added by refinement from 50 kb to 5 kb compartment mammoth vs. elephant as measured by the inverse of the autocovariance function. **(E)** 5 kb CRUSH scores in mammoth skin (x-axis) vs. ancient RNA (aRNA) levels from mammoth muscle (y-axis). **(F)** Similarity between 5 kb CRUSH calls in the mammoth skin vs. the skin, liver, ovary, PBMC, or brain of elephants. **(G)** 5 kb compartment tracks and the location of POU6F2 motifs near *EGFR*.
